# Comparative nectar metabolomics reveals sucrose-nitrogen tradeoffs and chemical drivers of microbial growth in floral nectar

**DOI:** 10.64898/2026.05.26.727296

**Authors:** Rachel L. Vannette, Caitlin Rering, Jacob M. Cecala, Leta Landucci, Alexia Lanier

## Abstract

**Introduction:** Many plant species secrete nectar to attract beneficial animals. The chemical composition of floral nectar influences pollinator nutrition and behavior, as well as microbial growth in flowers. Yet factors that predict nectar composition across plant species, as well as chemical compounds determining microbial growth in nectar, remain poorly understood.

**Methods:** We used both targeted and untargeted metabolomics to compare the nectar chemical profiles across 31 phylogenetically diverse plant species that span a range of floral morphologies. We examined the common classes of compounds detected in nectar and patterns of co-occurrence among them. We combined newly collected chemical data with previously published data on microbial growth in nectar of the same plant species to examine how nectar chemistry is associated with microbial growth.

**Results:** Plant species and clades varied in amino acid, minor sugar, and secondary metabolite composition and concentration. Sampled rosids and lilioids generally contained higher amino acids while asterids contained greater concentrations of oligosaccharides and sugar alcohols. Across plant species, proteinogenic amino acids frequently co-occurred in nectar but many were negatively associated with sucrose concentration. Plant species with greater concentrations of amino acids and other nitrogen-containing compounds hosted greater microbial density in nectar, while some other compound groups were negatively associated with microbial diversity.

**Conclusions:** Negative correlations between nectar amino acid and sucrose concentration across species suggest ecological tradeoffs or physiological constraints in nectar composition. Given that the growth of common nectar microbes is limited by amino acid concentration, these findings suggest an ecological cost to amino acid production in nectar. Finally, we document variation among species in nectar vitamins, non proteinogenic amino acids and secondary metabolites with hypothesized yet currently untested ecological roles.

## Introduction

Floral nectar serves as the primary reward in the pollination mutualism between angiosperms and animals. Nearly 75% of animal-pollinated plants, including over 220,000 angiosperm species, produce nectar (Ballarin *et al*., 2024), and many animal groups depend on nectar as a primary carbohydrate source. Nectar chemical composition is variable among plant species but is typically an aqueous solution of mono- and di-saccharides containing dilute concentrations of amino acids, proteins, ions, lipids and diverse plant metabolites (Baker, 1977; Nicolson *et al*., 2007; Roy *et al*., 2017; Palmer-Young *et al*., 2019a). Nectar is produced and secreted via specialized processes including enzymatic degradation of stored starch, biosynthesis of sucrose and amino acids, and transport of nectar constituents (Roy *et al*., 2017). Although nectar is generally thought to be tailored for pollinator attraction and reward (Baker, 1977), given the high metabolic and ecological costs associated with the production and maintenance of nectar (Southwick, 1984; Pyke, 1991), plants are likely under selection to defend nectar from exploitation by microbes and other antagonists via the evolution of chemical or other defenses (Adler, 2000; Antonovics, 2005; González-Teuber & Heil, 2009; McArt *et al*., 2014; Rivest & Forrest, 2024).

Nectar chemical composition affects nectar feeding organisms in ways that feed back to influence plant reproduction. For example, nectar sugar concentration and composition affects foraging of adult honey bees (Vansell, 1934), developmental success of larval solitary bees (Burkle & Irwin, 2009), and fitness of butterflies and parasitoids (Wäckers, 2001; Cahenzli & Erhardt, 2012). Secondary metabolites in consumed nectar can reduce the growth of bee parasites (Richardson *et al*., 2015; Koch *et al*., 2019), affect honey bee memory (Wright *et al*., 2013) or enforce pollinator specialization (Tiedeken *et al*., 2016; Cane *et al*., 2020). In addition, metabolites that contribute pigmentation to nectar may act as foraging signals for floral visitors or antimicrobial properties (Hansen *et al*., 2007; Magner *et al*., 2024). However, metabolite effects on pollinators may depend on additional floral cues (Jones & Agrawal, 2022) or the composition of chemical mixtures (Muth *et al*., 2022), while metabolites themselves are likely to be multifunctional (Junker & Blüthgen, 2010). As a result, the composition of low-abundance compounds in nectar is of increasing interest due to their demonstrated or hypothesized influence on pollinators (Stevenson *et al*., 2017; Nicolson, 2022; Barberis *et al*., 2023; Hemingway *et al*., 2024) and subsequent effects on plant fitness.

Nectar metabolites may also influence nectar exploitation by fungi and bacteria (González-Teuber & Heil, 2009). Microbes found in nectar include plant pathogens (Antonovics, 2005; McArt *et al*., 2014; Vannette, 2020) or nonpathogenic microbes which can affect pollinator visitation (Herrera *et al*., 2008; Vannette *et al*., 2013; Schaeffer *et al*., 2017; Yang *et al*., 2019) or reduce pathogen growth (Rering *et al*., 2023). Microbial selection may act on floral traits (Rebolleda-Gómez *et al*., 2019) yet it remains unclear what nectar traits are important in shaping microbial abundance and composition (Schmitt *et al*., 2021; Francis *et al*., 2023; Cecala *et al*., 2025). Although individual compounds can influence microbial growth in nectar and nectar-like solutions (reviewed by Schmitt *et al*. 2021), how variation in nectar composition across plant species shapes microbial abundance and composition in nectar has not previously been examined in a comparative framework. We hypothesize that chemical resource availability, either amino acid concentration or vitamin concentration, may limit microbial growth (Christensen *et al*., 2021; Chappell *et al*., 2024). Although microbial growth in nectar is likely not sugar-limited, high sugar concentration and resulting osmotic stress may reduce microbial growth (Herrera *et al*., 2009a). On the other hand, previous work suggests that some metabolites including alkaloids (Wink, 2008), terpenoids (Whitehead, 1999) or phenolics may directly limit microbial growth in nectar (Burdon *et al*., 2018; Boachon *et al*., 2019; Mueller *et al*., 2023).

Although many adaptive hypotheses for nectar chemical composition including pollinator manipulation and antimicrobial effects have been proposed, some nectar constituents may be the product of expression or biosynthesis in other plant tissues (Adler, 2000) and result from nonspecific transport, or absorption or solubility in aqueous solutions (Raguso, 2004). Indeed, defensive chemistry in leaf tissue is predictive of nectar chemistry in the genera *Nicotiana* (Adler *et al*., 2012), however, neither *Asclepias* nor *Salvia* show strong correspondence between leaf and nectar metabolites (Manson *et al*., 2012; MacNeill *et al*., 2025). Because many secondary compounds are restricted to particular plant clades or ecological or defense strategy (Wink, 2003), we expect that the composition of some nectar compounds may reflect phylogenetic conservatism among plant clades.

To survey and explore the consequences of variation in nectar composition across plant species, we use both targeted and untargeted metabolomic analyses to characterize the chemical composition of floral nectar in 31 phylogenetically diverse plant species. We pair these metabolomic data with a recently published dataset examining growth of inoculated microbes in the nectar of the same plant species (Cecala *et al*., 2025). We leverage these datasets to ask the following questions: 1) What are the common classes of compounds detected in nectar, and do plant clades or species vary in nectar chemical composition? We first assessed chemical composition broadly using untargeted LC-MS/MS, then used a targeted approach to quantify sugars, amino acids and vitamins known to be important to floral visitor preference and performance. With these data, we examined: 2) What compounds frequently co-occur in floral nectar across plant species? 3) Does plant pollination guild or phylogenetic relatedness explain variation in nectar chemical composition? 4) What components of nectar chemistry predict variation in microbial growth or community composition in floral nectar?

## Methods

### Sample collection

We sampled nectar from 31 phylogenetically diverse plant species, spanning monocots and dicots, and including 20 plant families (Table S1). All plant individuals were growing on the UC Davis campus or Sagehen Reserve and sampled in 2023. Nectar samples were collected from flowers that had been bagged with a mesh enclosure before anthesis and allowed to open in the bag, to increase nectar accumulation and exclude pollinators and the microbes that they vector (see Cecala *et al*. 2025 for experimental details). Flowers were excised from the plant in the field and transported back to the lab in clean tip boxes where nectar was extracted within 1 hour of collection. Nectar was collected using clean capillary tubes either via the top or bottom of the corolla, and care was taken to remove anthers or minimize pollen contamination. For most plant species, nectar samples were pooled from multiple flowers on the same plant, and accumulated volumes ranged from 5 µl to over 100 µl. Plant species were sampled on multiple days (Cecala *et al*. 2025, supporting information) and when available, replicates were chosen from different sampling dates and different plant individuals. Nectar samples were stored at -80 °C or on dry ice until analysis. Three samples had insufficient volumes to allow the full suite of chemical analyses and were only included in untargeted metabolomics and targeted amino acid analysis. Blanks (n=8) were created by pipetting sterile water using capillary tubes and processed in the same way as nectar samples above.

### Sample processing and chemical analysis: untargeted and targeted metabolomic analyses

#### Sample preparation

Samples and method blanks were diluted with 70% methanol (Optima™, Fisher Scientific, Waltham, MA, United States) and filtered (0.22 µm pore, 4 mm diameter, Millex™ PVDF, Millipore, Burlingon, MA, United States). Three dilution levels were prepared serially based on the recorded nectar volume: 25-fold for nontargeted chemical analysis and targeted amino acid and pantothenic acid analysis, 250-fold for minor sugar analysis, and 2500-fold for major sugar analysis (fructose, glucose, and sucrose). Pooled samples and blanks were separately prepared from each of the three dilution levels. Pooled nectar samples and blanks were prepared by combining aliquots of each sample and method blank, respectively. The pooled sample served as a quality control (QC) technical replicate injected after every 9 samples, and the pooled sample and pooled method blank formed the basis for data-dependent MS^2^ generation.

### Untargeted metabolomics

Samples were analyzed by liquid chromatography mass spectrometry (LC-MS). Full method details are provided in the supplement (Supplementary methods S1). Briefly, samples were analyzed with a Hypersil GOLD™ C18 column (2.1 × 150 mm, 1.9 µm, Thermo Scientific, Waltham, MA, United States) and a binary mobile phase system of 0.1% formic acid in water and acetonitrile. LC eluate was directed to an IQ-X Orbitrap MS. All samples and blanks were analyzed in positive and negative full scan profile mode (MS^1^; 50-800 m/z). The pooled method blank and pooled sample were analyzed with the AcquireX platform to collect data-dependent MS^2^ spectra. In this process, exclusion and inclusion lists were first generated via MS^1^ analysis of the pooled method blank and pooled sample, respectively. Then, iterative MS^2^ analyses of the pooled sample were performed. In our MS^2^ AcquireX-enabled method, [M+H]^+1^ and [M-H]^-1^ were selected as preferred precursors and were fragmented by higher-energy collisional dissociation (HCD) and by collision-induced dissociation (CID). Product ions from both HCD and CID fragmentation were detected in profile mode by the orbitrap at 30,000 resolution with auto scan range mode enabled.

### Data processing and feature annotation

Full data processing and feature annotation details are provided in Supplementary methods S2. Data were centroided and exported to .mzML format for processing using MZmine software version 4.1.0 (Schmid *et al*., 2023). Spectra were processed by the Mass detection module and features detected using the Chromatogram builder (Myers *et al*., 2017). Overlapping features were resolved using the Local minimum resolver algorithm and Isotope pattern finder. Features were then aligned across samples with Join aligner, filtered with the Feature list rows filter, and grouped with Correlation grouping (i.e., metaCorrelate) according to feature shape. Ion identity molecular networking (IIMN) was used to additionally group features that arose from the generation of multiple ion species of the same compound (e.g., [M+H]+ and [M+Na]+; Schmid *et al*., 2021). We used the Export molecular networking files module to generate files (MS^2^ MGF files, feature quantification table with areas, and ion identity edges from IIMN) for feature based molecular networking (FBMN, release 28.2, (Nothias *et al*., 2020)) via the Global Natural Products Social molecular networking platform (GNPS, (Wang *et al*., 2016)). Retained features were annotated using SIRIUS v6.0.5 (Dührkop *et al*., 2019) relying on CANOPUS (Dührkop *et al*., 2021), Classyfire and CSI:FingerID. Annotations were combined into a feature table for downstream analyses. Untargeted compound diversity within a sample was estimated using Hill number N=1 (Chao *et al*., 2014).

### Targeted metabolomics

#### Carbohydrates

Nectar carbohydrates were quantified via targeted LC-MS analysis. Full analytical details are provided in Supplementary methods S3. Sugars were separated on a Hypersil GOLD™ Amino column (100 × 2.1 mm, 1.9 μm, Thermo Scientific) and isocratic 2 mM ammonium acetate in 80:20 acetonitrile/water adjusted to pH = 5.4 with acetic acid at 0.5 ml min^-1^. The MS was operated in single reaction monitoring mode. For major sugars (glucose, fructose, and sucrose), a 3 min run time was used and samples were injected after 2500-fold dilution. Minor sugar components (myo-inositol, erythritol, maltose, maltotriose, mannitol/sorbitol (measured as one feature due to co-elution and shared precursor and product ions), melezitose, melibiose, raffinose, ribose, stachyose, trehalose, xylitol) had a run time of 16 min and 250-fold dilution. Sugars were quantified with external calibration standards.

#### Amino acid and pantothenic acid quantification

Preliminary data analysis via Compound Discoverer (v. 3.3, Thermo Fisher Scientific) tentatively identified 21 amino acids (Supplementary Methods, Table S2) and the vitamin pantothenic acid in samples. We injected commercial standards to validate compound identities according to retention time, [M+H]+ precursor ion exact mass, and CID and HCD fragmentation patterns. External calibration standards were used for post-hoc quantification of these metabolites from MS^1^ positive mode data.

### Microbial growth data

We used data from a previous publication to assess microbial growth potential in floral nectar across 28 of the 31 plant species. We inoculated a fixed microbial community and examined establishment after 24 hours to assess microbial growth and community assembly in flowers where pollinators were excluded. For full methods on microbial synthetic community preparation and floral inoculation see Cecala *et al*. 2025. Briefly, during the spring of 2023 we inoculated previously unopened and bagged flowers with a synthetic community comprised of two common yeast species (*Metschnikowia reukaufii* and *Aureobasidium pullulans*) and three bacterial species (*Acinetobacter pollinis*, *Apilactobacillus micheneri*, and *Neokomagataea thailandica*). After 24 hours of growth, flowers were harvested, nectar collected and plated on selective media to quantify the abundance (colony forming units) of each focal microbe. Flower bagging was largely successful in excluding floral visitation except for rare occurrences of ants within bags. Resulting microbial abundance, averaged by plant species, ranged over three orders of magnitude, from 9.0±6.8 µL^-1^ in *Arbutus unedo* to 3.2±1.7 × 10^4^ µL^-1^ in *Hesperaloe parviflora*.

### Statistical analyses

#### 1) What are the common classes of compounds detected in floral nectar across plant species?

To visualize the common classes of compounds detected in the untargeted analyses across plant species and their structural similarity, we used Cytoscape to generate feature-based molecular network of compounds, including all detected compounds. We then subset to only compounds annotated with 75 or greater confidence (Fig S1) and used only annotated compounds for further analysis, focusing on data acquired in positive mode in the main text (but see supplementary tables for negative mode data). To compare if compound diversity in the untargeted dataset differed among plant species or clades, we compared observed compound richness among species using linear models. To compare if compound composition differed among plant species, we used PerMANOVA based on Jaccard distance with plant clade (rosids, asterids or lilioids) plant species as a predictor and F-tests to estimate significance. For subsequent analyses, we calculated species mean values.

We examined if plant species or clades varied in the composition of targeted analytes using nonmetric multidimensional scaling and permutational ANOVA (PerMANOVA) using Bray-Curtis dissimilarities and examined compounds that distinguished these groups using envfit implemented in the R package vegan (Oksanen *et al*., 2007). We used linear models to examine if targeted analytes differed in concentration among plant species or clades, with separate models for each.

#### 2) What compounds frequently co-occur in floral nectar across plant species?

To examine potential covariation among compound classes (untargeted dataset) or quantified metabolites (targeted dataset), we used Pearson correlations for individual metabolites from the targeted dataset or summed compound abundances within each NP Classifier superclass, each averaged at the plant species level. To compare between results from the targeted and untargeted analyses, we used species mean values to assess correlations among chemical classes.

#### 3) Does plant pollination guild or phylogenetic relatedness explain variation in nectar chemical composition?

To assess the contribution of floral morphology, we scored floral phenotypes of all plant species on the basis of 28 binary traits used in past studies (Faegri & Pijl, 1979; Ollerton *et al*., 2009) to represent pollination syndromes in multivariate space using Bray-Curtis dissimilarity. Hierarchical cluster analysis was used to classify plant species into two guilds, one with traits consistent with bird pollination and the other with bee or other insect pollination. For full details, see Cecala et al. (2025). We used PerMANOVA to examine if pollination syndrome and plant clade was associated with variation in targeted or untargeted metabolite composition using Bray-Curtis and Jaccard dissimilarities, respectively.

We estimated plant phylogenetic relationships among sampled plant species using the function ‘phylo.maker’ in package *V.PhyloMaker2* (Jin & Qian, 2022) using the reference vascular plant mega-phylogeny GBOTB.extended.TPL. To test for relationships between plant phylogenetic relatedness and multivariate compound composition, we created a pairwise distance matrix of plant phylogenetic relatedness using function ‘cophenetic.phylo’ in package *ape* (Paradis & Schliep, 2019). We compared this distance matrix to a dissimilarity matrix of the untargeted compound composition (Jaccard dissimilarity) or the abundance of sugars or amino acids in the targeted analysis (Bray-Curtis dissimilarity) using a Mantel test via function ‘mantel’ in package *vegan*, calculating Spearman’s ρ with 10,000 permutations.

### 4) Is microbial growth or composition in floral nectar associated with nectar chemistry?

We used species-level average compound composition from the untargeted metabolomics dataset (positive mode) and the targeted analyses to examine associations with microbial abundance and composition in flowers of that species inoculated with a synthetic community. Here, nectar chemical composition represents conditions found in uninoculated flowers, the conditions encountered upon microbial arrival to a flower rather than following microbial growth.

Within the subset of plant species where microbial growth was assessed (N=28 species), we examined first if metabolite composition was associated with total microbial growth or Shannon diversity using separate PerMANOVAs for targeted and untargeted analysis using Bray-Curtis or Jaccard dissimilarity matrices respectively. Second, we examined if metabolite composition was associated with variation in microbial composition as Jaccard distance using Mantel tests. Third, we used multiple regression to examine if hypothesized predictors including the total concentration of amino acids, vitamin B5 (pantothenic acid), or sugar concentration, was associated with either total microbial density or Shannon diversity. We used additional Mantel tests to examine the associations between plant phylogenetic distance, pollinator syndrome and metabolite dissimilarity.

Finally to examine if particular groups of metabolites were associated with variation in microbial growth, we used co-expression patterns to generate topological overlap, which were used to generate modules (min 10 compounds), using package WGCNA (Langfelder & Horvath, 2008). This approach reduces the number of independent variables (chemical compounds) and spurious correlations while accounting for correlational expression of compounds in metabolomics datasets (Forister *et al*., 2020). Compound modules were examined for correlations with microbial density in nectar, Shannon diversity or nectar volume.

Figures were created using package *ggplot2* (Wickham, 2016) and tree plots using ggtree (Yu *et al*. 2017) and custom function ‘ggtreeplot’ (Hackl 2018). Data wrangling was performed using phyloseq and microViz (McMurdie & Holmes, 2013).

## Results

### 1) What are the common classes of compounds detected in floral nectar across plant species?

#### Untargeted analytes

After processing with MZmine and blank subtraction, 3168 features were identified in the untargeted metabolomics dataset in positive mode and 1243 in negative mode across all nectar samples considered. Library and in-silico based classification of fragmentation spectra and parent ions via GNPS and SIRIUS generated annotation at 75% confidence for 352 features in positive mode (Table 1, Table S1), leaving 2816 features (∼88%) unannotated. We restrict further analyses to annotated features which we refer to as ‘compounds’ and focus on compounds detected in positive mode (see Tables S2-3 for negative mode results).

**Table 1.** Classification of annotated compounds detected in positive mode using NP Classifier as implemented in CANOPUS and SIRIUS. Feature richness is across all samples in the dataset after blank subtraction.

| Natural Products<br>Classification<br>Pathway | Rich<br>ness | Example<br>structures | Class | Hypothesized ecological roles |
| --- | --- | --- | --- | --- |
| Alkaloids | 52 | Purine alkaloids, Polyamines, Imidazole alkaloids, Pyridine alkaloids, Piperidine alkaloids, Pyrrolidine alkaloids, Acetate-derived alkaloids, Carboline alkaloids, Simple indole alkaloids, Pyrrolizidine alkaloids, Isoquinoline alkaloids, Phenylalanine-derived alkaloids, Quinoline alkaloids, Quinolizidine alkaloids, Anthranillic acid derivatives, Steroidal alkaloids |  | Deterrence, antimicrobial activity |
| Amino acids and<br>Peptides | 61 | Amino acids, Tripeptides, Cyanogenic glycosides, Dipeptides, Cephalosporins, Linear peptides, Aminoglycosides, Microcolins and mirabimids |  | Pollinator attraction, primary<br>metabolites and metabolic<br>byproducts or defense compounds |
| Carbohydrates | 58 | Polysaccharides, Monosaccharides, Disaccharides, Aminosugars, Cyclitols, Shikimic acids and derivatives, Purine nucleosides, Cyanogenic glycosides, Purine alkaloids, Amino acids, Aminoglycosides |  | Pollinator reward, primary<br>metabolites |
| Fatty acids | 122 | Halogenated fatty acids, Sphingoid bases, Fatty alcohols, Fatty aldehydes, Unsaturated fatty acids, Primary amides, Monosaccharides, Amino acids, Dicarboxylic acids, Oxo fatty acids, Hydrocarbons, Thia fatty acids, Fatty acyl homoserine lactones, Fatty acyl carnitines, N-acyl amines, Oxygenated hydrocarbons, Jasmonic acids, Other Octadecanoids, Phosphosphingolipids, Glycerophosphocholines, Nitro fatty acids, Amino fatty acids, Monoacylglycerols, Ceramides |  | Primary metabolites, hypothesized<br>roles in evaporation reduction,<br>pollinator attraction, signaling |
| Polyketides | 10 | 2-pyrone derivatives, Naphthoquinones |  | Hypothesized antimicrobial activity |
| Shikimates and<br>Phenylpropanoids | 27 | Simple phenolic acids, Cinnamic acids and derivatives, Simple coumarins, Flavones, Shikimic acids and derivatives, Kavalactones and derivatives, Gallotannins, Flavonols, Flavanones |  | Pigmentation/coloration, antioxidant<br>activity, possible antiherbivore<br>activity |
| Terpenoids | 22 | Iridoids monoterpenoids, Secoiridoid monoterpenoids, Menthane monoterpenoids, Apocarotenoids(ε-), Bisabolane sesquiterpenoids, Monocyclic monoterpenoids, Eudesmane sesquiterpenoids, Androstane steroids, Thia fatty acids, Chalcones |  | Signaling, hypothesized deterrence or<br>antimicrobial activity |

Using untargeted positive mode data, plant species ranged from 70-166 annotated compounds detected with an average of 106 detected (Fig 1) and differed in the number of compounds observed (F_31,60_=5.38, p<0.001). Plant species identity explained 87% of variation in untargeted metabolite composition among samples (PerMANOVA p<0.001; Fig 1). The greatest number of compounds were annotated within the fatty acids (122), amino acids and peptides (61), carbohydrates (58) and alkaloids (52) pathways; Table 1. Feature-based molecular networking connected compounds assigned to shared and differing pathways (Fig S1), suggesting common and cascading biosynthetic pathways are active in nectar/nectaries. Many carbohydrates, a core set of fatty acids, pyrone derivative polyketides, and some phenolic acids were detected in nearly all plant species (Fig 1) while terpenoids, alkaloids, and a subset of fatty acids were more sporadically detected, with compound subclasses detected only in a subset of plant species (Fig 1A).

**Figure 1.**
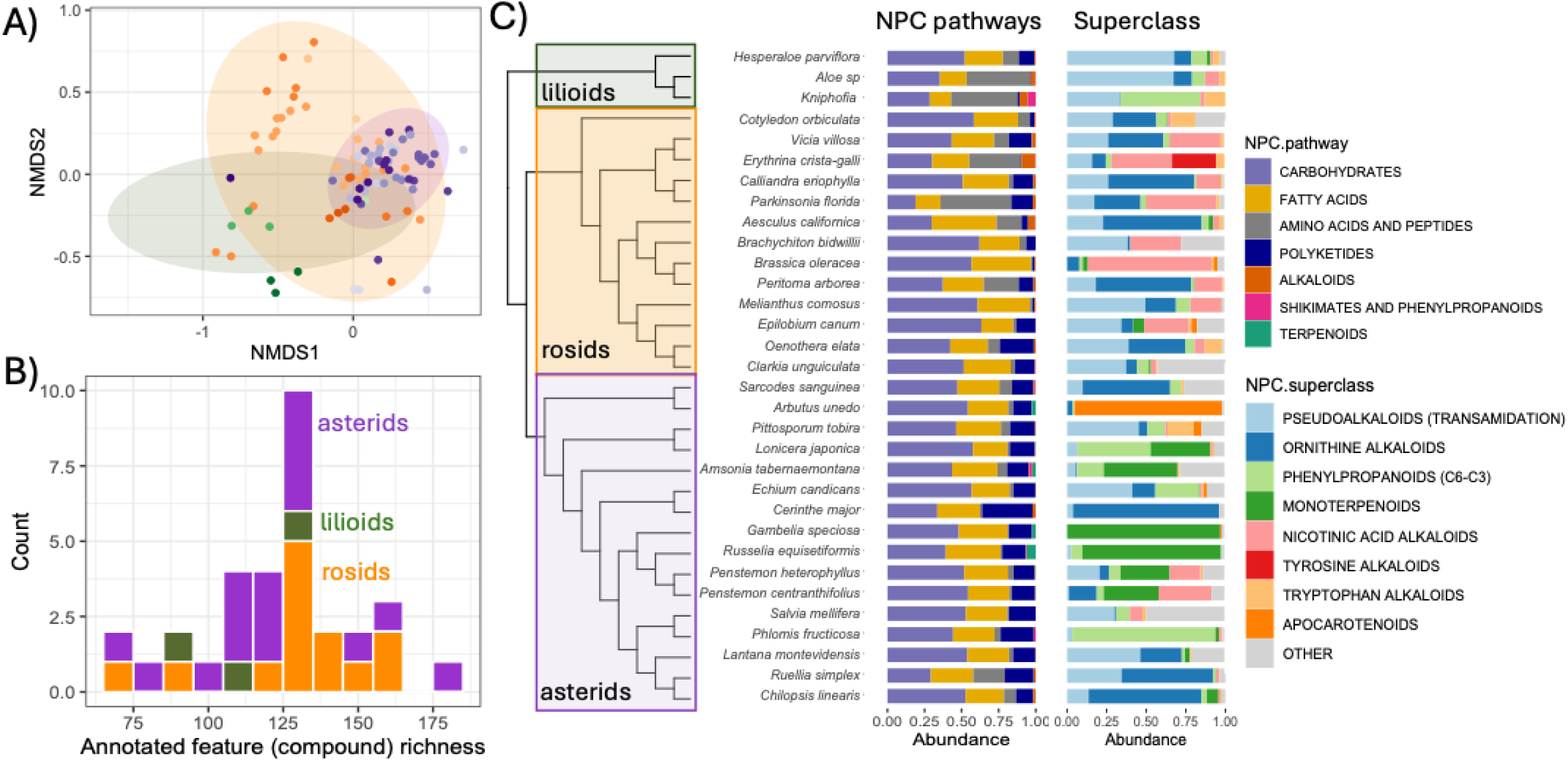
Untargeted metabolite composition and diversity differ among plant species and clades. In A) the composition of untargeted analytes varies among plant species (R^2^=0.75; p<0.001) and clades (R^2^=0.05; p<0.001) using NMDS based on Jaccard dissimilarity of all annotated compounds, with stress=0.2. Points indicate individual samples and color families indicate the clade of the plant species with green indicating lilioids, orange indicating rosids and purple for asterids. Shaded ovals indicate 95% CI. In B) observed compound diversity of annotated chemicals varies among plant species (F_29,89_=6, p<0.001) but not clade (F_2, 89_=0.5, p=0.57), with the count of plant species with a given annotated feature richness. In C) phylogeny of sampled plant species colored by plant clade and corresponding averaged compositional barplots display the relative abundance of Natural Product Category pathways (first column) and secondary metabolite superclasses within alkaloids, shikimates and phenylpropanoids, and terpenoids (second column). We note that untargeted data collected are not strictly quantitative. Colors indicate plant clade across panels.

Metabolites frequently detected in nectar in the untargeted dataset included sugars, amino acids and fatty acids (Table 1), but also the phenylpropanoids coumaric acid and homogentisate, which both exhibit antioxidant and antimicrobial activity. Both phenylpropanoids are essential metabolic precursors, and were detected in nearly all plant species. In addition, the terpenoid abscisic acid (i.e., abscisin), a plant hormone controlling ripening and floral development was detected in more than half of examined species. Additional carbohydrates detected widely included levoglucosan (i.e., glucosan), galactosucrose (i.e., lactosucrose), isomaltose (i.e., brachiose), and umbelliferose. Other compounds were restricted to specific species or lineages including the iridoid glycosides geniposidic acid, catalpol, aucubin, as well as syringin and the insect antifeedant pterocarpol. Many fatty acids were widely detected including carboxylic acids and their derivatives, carbonyls as well as butanolide (a water soluble mycotoxin), citrate and choline. Notably, some individual plant species hosted higher diversity and abundance of phenolics (e.g. *Lonicera*, *Cerinthe*, *Melianthus* among others) or alkaloids (e.g. the legumes *Erythrina* and *Parkinsonia*).

#### Targeted analytes

Our targeted analysis focused on 15 sugars, 21 amino acids, and vitamin B5 (pantothenic acid). Plant species differed significantly in the composition of amino acids, and major and minor sugars (Fig 2), with plant species identity explaining between 63-82% of variation in targeted metabolite composition. In addition, plant clades differed generally in the composition of major sugars, amino acids and minor sugars/sugar alcohols (Fig 2). Lilioids (monocots) contained the lowest average sucrose and minor sugar concentration (F_2,89_=7.8, p=0.0007), nearly 50% less than rosids or asterids (Fig S2-S3). In contrast, lilioid nectar contained the highest amino acid concentration (F_2,89_=10.3, p<0.0001), more than 10 times greater than average in the asterids and 2.5 times greater than the rosids. The signaling amino acid AABA was frequently detected in rosids but not in asterid species examined here, while the sugar alcohols myo-inositol, trehalose and xylitol were frequently detected in rosids and asterids but rarely detected in sampled lilioids.

**Figure 2.**
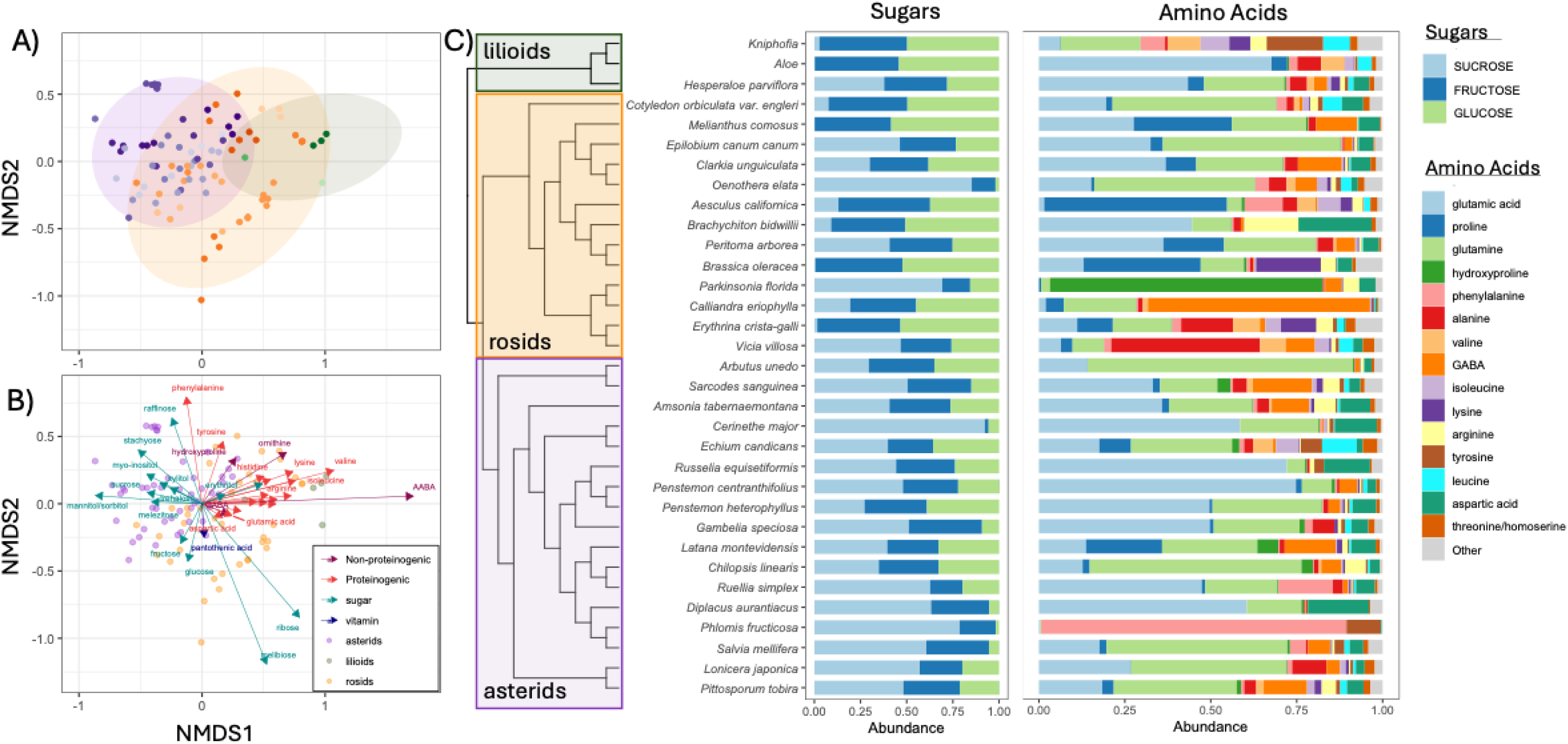
Targeted metabolites (amino acids, sugars, sugar alcohols, pantothenic acid) differ in composition and concentration across plant clades and species. A) Unconstrained ordination based on Bray-Curtis dissimilarities using nonmetric multidimensional scaling. Points indicate individual nectar samples colored by plant species, with color families organized by clade. In B) the same ordination is shown with analyte classes superimposed and colored by type. In C) the relative abundance of sugars (first column) and amino acids (second column) are shown. Colors indicate plant clade as in Fig 1.

### 2) What compounds frequently co-occur in floral nectar across plant species?

We detected covariation among compound abundance both in our targeted analysis (Fig 3a) as well as among superclasses detected in the untargeted metabolomics dataset (Fig 3b). We noted positive associations among many amino acids, with proteinogenic amino acids often co-occurring in nectar (Fig 3a). In contrast, higher sucrose concentration was negatively associated with the concentration of glucose and many amino acids. In the untargeted dataset, clusters of correlated compounds included polyols, ornithine alkaloids, fatty acid conjugates, coumarins and carotenoids (Fig 3b). Other compound classes also co-occurred including tryptophan alkaloids, phenylpropanoids, pseudoalkaloids, small peptides and tyrosine and nicotinic acid alkaloids.

**Figure 3.**
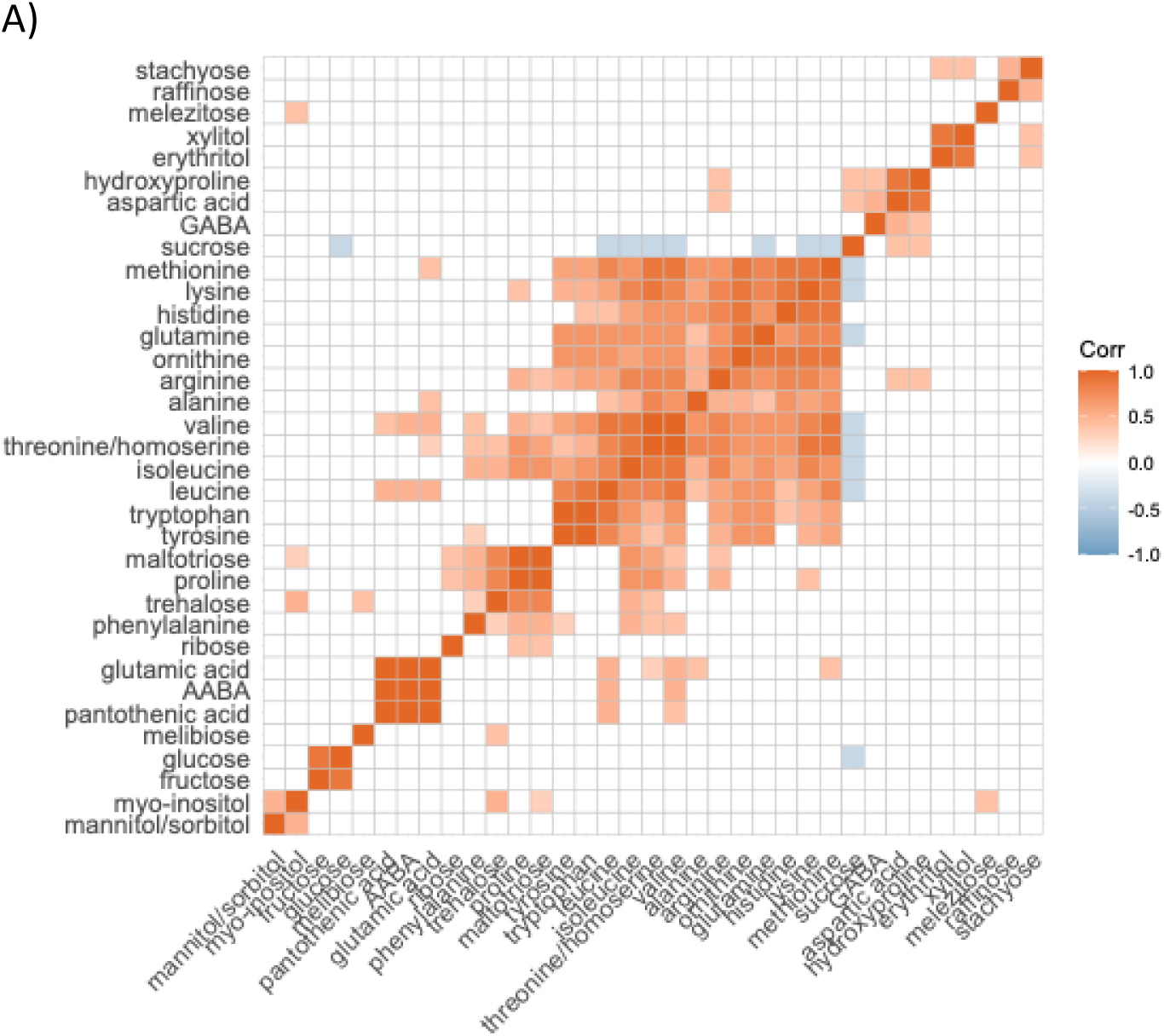
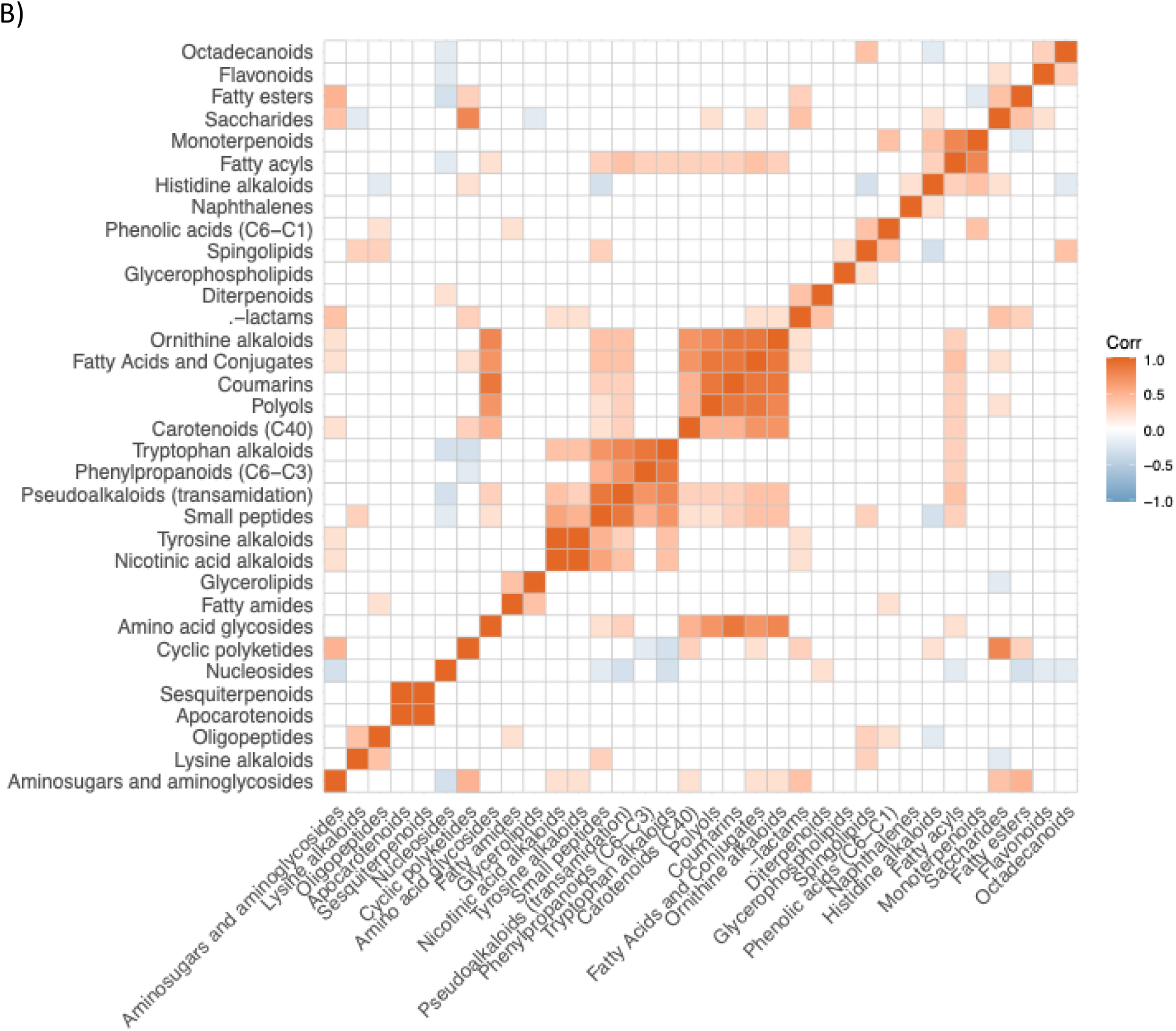
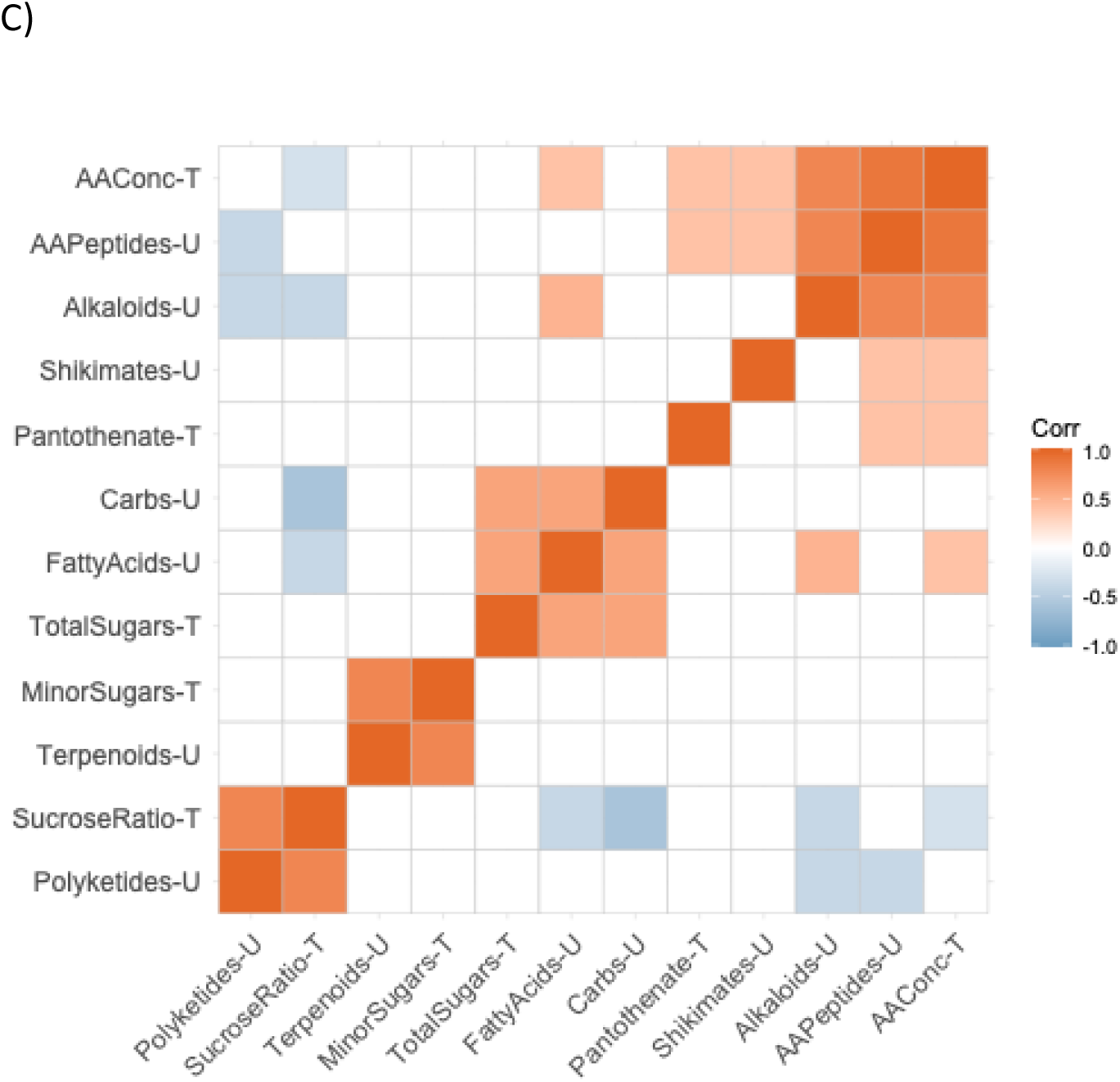
Covariation in species means of A) sugars and amino acids detected in the targeted metabolite dataset and B) compound superclasses assigned to untargeted metabolomic compound matrix among species. The strength of Pearson correlations is shown among species averages. C) shows correlations between metabolites in the targeted (T) and untargeted (U) datasets. In all three, correlations are indicated in squares with red indicating positive correlations and blue indicating negative associations, with correlations p>0.05 not shown. Compounds are ordered according to their similarity based on hierarchical clustering.

Across species, alkaloids, amino acids and fatty acids were positively associated, while these were negatively associated with polyketide and sucrose concentrations. Shikimates were positively associated with total amino acid concentration, and terpenoids were positively associated with minor sugar concentration (Fig 3c).

### 3) Does plant pollination ecology or phylogenetic relatedness explain variation in nectar chemical composition?

We used floral traits to designate each species’ pollination ecology (Cecala et al. 2025) which in previous analyses was best represented by two clusters: either plant species with red flowers, abundant nectar and long corollas, associated with bird pollination (Cluster 1) or shorter flowers, less nectar and open corollas, associated with bee pollination (Cluster 2). Floral morphology was associated with nectar composition: nectar from Cluster 2 flowers had at least 1.5 times the concentration of sucrose, glucose and fructose than did cluster 1 flowers (p<0.01 for all). Cluster 2 nectar contained approximately 1.5 times greater concentrations of minor sugars and sugar alcohols compared to those in Cluster 1 (F_1,76_=22.99, p<0.001). Cluster 2 also contained higher average amino acid content but this was in large part driven by the presence of lilioids within this category and not significant when they were removed. Plant species pollination syndrome (summarized by a matrix of floral traits) was also associated with variation in sugar composition (Mantel r =0.11, p=0.03) and amino acid composition (Mantel r=0.12, p=0.02). Pollination clusters differed in the composition of untargeted metabolites (Fig 4, PerMANOVA F_1,79_=3.91, p<0.001). In addition, 36 compounds from multiple pathways were differentially abundant between pollination clusters at FDR<0.001 (Fig 4b).

**Figure 4.**
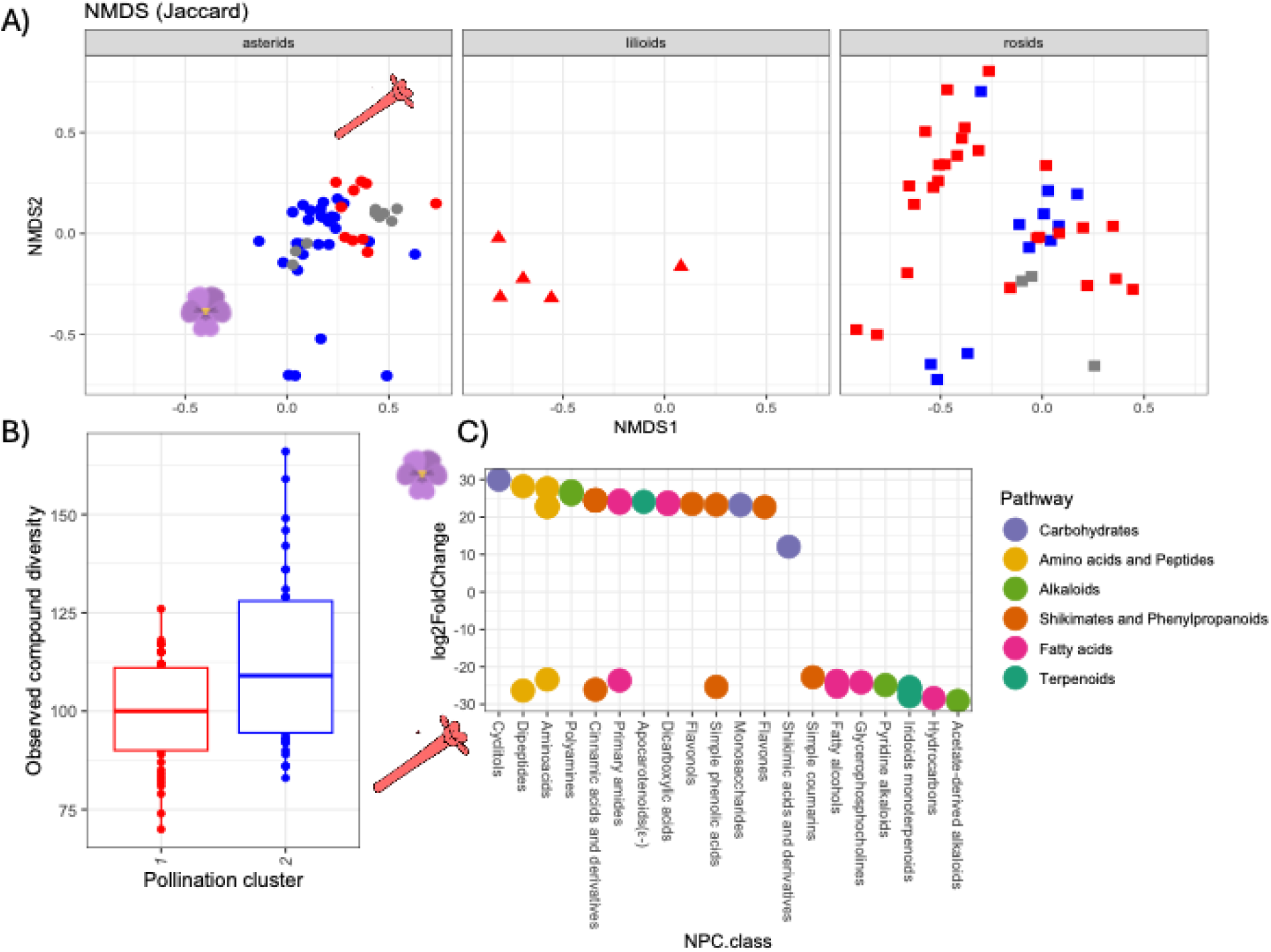
Untargeted metabolites vary based on pollination cluster A) based on NMDS ordination, with red indicating traits typically associated with bird pollination and blue indicating traits associated with bee pollination. Points represent individual nectar samples, and faceted by plant clade. NMDS was performed with all data points and are facetted for visual interpretation. B) Compound diversity differs between pollination clusters and C) Compounds differentially abundant between pollination clusters, with values below 0 indicating greater abundance in cluster 1 (bird pollination) and above 0 indicating greater abundance in cluster 2 (bee pollination) using DESeq with FDR<0.001.

Across plant species, increasing phylogenetic distance was associated with species’ dissimilarity in untargeted metabolite composition (Fig 1, Mantel r=0.29, p<0.001, Fig S4-S8), major sugars (Mantel r=0.29, P<0.001) whereas the phylogenetic distance was marginally associated with amino acid composition (Mantel r=0.13, p=0.06).

### 4) Is microbial growth or composition in floral nectar associated with nectar chemistry? Nectar traits predict variation in microbial growth

Across plant species, untargeted metabolite composition was associated with variation in microbial diversity maintained in plant nectar (PerMANOVA R^2^=0.11, p=0.026), but was not significantly associated with variation in microbial abundance (PerMANOVA R^2^=0.07 p=0.14). In addition, compound diversity in the untargeted dataset was not associated with variation in microbial abundance or diversity (p>0.10).

As predicted by resource limitation hypotheses, a plant species’ average total concentration of free amino acids in nectar was positively associated with microbial abundance in nectar (t=2.4, p=0.02; Fig 5). In addition, microbial diversity decreased with increasing free amino acids in nectar (t=7.14, p=0.01, Fig 5b), suggesting that competitive dominance by microbial species may be enhanced by high nitrogen availability within floral nectar. In contrast, neither the concentration of major sugars (sucrose, glucose and fructose) nor minor sugars and sugar alcohols was associated with microbial growth nor microbial diversity (p>0.4 for both).

**Figure 5.**
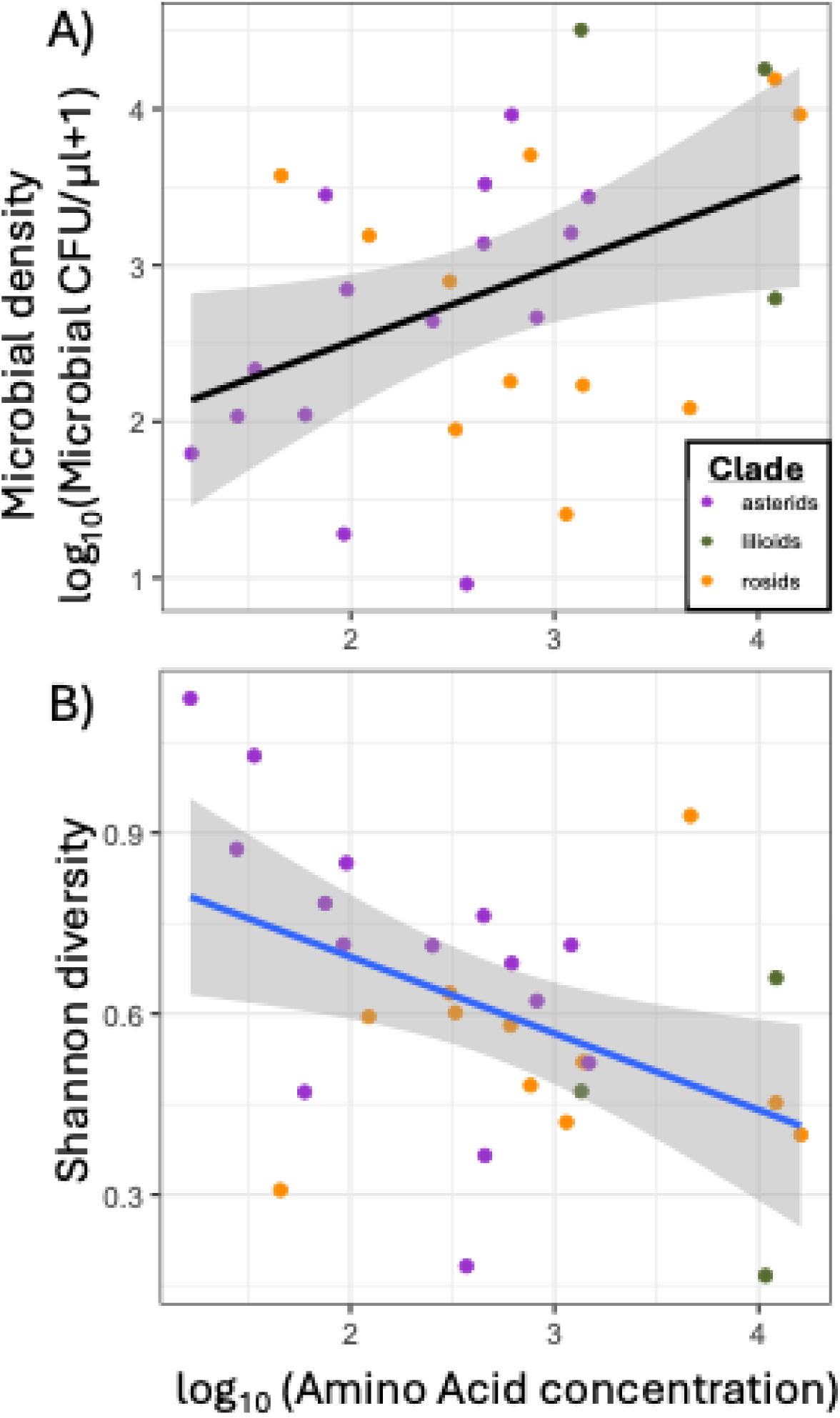
Amino acid concentration in floral nectar is positively associated with A) microbial density and B) microbial diversity in nectar among plant species. Each point represents the mean value for a given plant species, with point color indicating plant clade.

Using compound co-expression analysis, 12 chemical modules within the untargeted dataset were detected (Fig S9). Of these, two were positively associated with microbial density (Fig S10, black, yellow, both p-adj<0.05), while one was positively associated with microbial diversity in nectar (green, p-adj<0.05). Compounds in the black module, associated with higher microbial abundance, included amino acids and acetate-derived alkaloids (Table S4); while the yellow module contained amino acids, pyridine alkaloids and fatty acids. Compounds in the green module, associated with greater microbial diversity, included polysaccharides, imidazole alkaloids, fatty aldehydes, 2-pyrone derivatives and cyanogenic glycosides.

## Discussion

Our study adds to the growing understanding of the chemical complexity of floral nectar across plant species as well as correlates of this variation (Palmer-Young *et al*., 2019b; MacNeill *et al*., 2025). By quantifying nectar sugars and amino acids and assessing their variation with additional metabolites across a broad phylogenetic range, our study provides unique insights by linking sources of variation and co-variation in compound occurrence and abundance in nectar among plant species to resultant microbial growth. In addition to major sugars and amino acids, our analyses reveal that some minor constituents and natural products superclasses are abundant and in some cases ubiquitous in nectar across the sampled species (Figs 1, 3).

Specifically, short-chain fatty acids including fatty acyls, amides, esters and sphingolipids were consistently detected across plant species. Fatty acids have been detected in nectar in some species previously (Baker, 1977; Kram *et al*., 2008; Bender *et al*., 2012) including beyond their predicted solubilities in aqueous solutions (Kram *et al*., 2008). Lipids may serve as rewards for pollinators (Lepage & Boch, 1968; Levin *et al*., 2017), although roles in evaporation or plant signaling may be hypothesized. Nitrogen-containing compounds including amino acids and alkaloids were also common, particularly in plant species from the Fabaceae which each hosted alkaloid classes that were unique among the plant species sampled here. Notably *Erythrina crista-galli* was enriched in quinoline and the indole alkaloids norboldine and erythraline, both recognized for their anti-inflammatory and therapeutic effects. Nectar alkaloids can be directly toxic to pollinators (Cane *et al*., 2020) or at some concentrations may reduce pathogen loads in bees (Manson *et al*., 2010; Richardson *et al*., 2015). Recent work describing the nectar metabolome within the genus *Salvia* showed that bird-pollinated plant species showed distinct alkaloid composition compared to bee-pollinated plant species (MacNeill *et al*., 2025). Our comparison of metabolites based on pollination clusters similarly revealed differentiation in pyridine and acetate-derived alkaloids among other compound classes (Fig 4b). We also note the frequent detection of compounds with antioxidant activity, including apocarotenoids, cinnamic acid derivatives, gallotannins and flavonols (Table 1), also noted in previous studies of nectar (Carter & Thornburg, 2004; Palmer-Young *et al*., 2019b). These compounds likely confer protective absorption of UV radiation or antioxidant activity against radicals generated in floral nectar, either through abiotic mechanisms or via the plant’s own antimicrobial defenses (Carter & Thornburg, 2004).

Notably, our analyses reveal that plant clades can differ substantively in composition and concentration of primary metabolites, as well as composition of secondary metabolites. The sampled asterids were often low in amino acids yet high in sucrose and minor sugar constituents; rosids contained higher concentrations of glucose and fructose, and more frequent detection of ribose, melibiose and maltotriose. The lilioids sampled here, albeit only three species sampled and all within the Asparagales, contained the highest amino acid concentrations, across nearly all proteinogenic amino acids (Fig S2). Deeper taxon sampling within all clades (but particularly within the monocots) would further distinguish between potential drivers of variation in nectar chemistry, including pollination ecology, climatic drivers and phylogeny. Yet these trends suggest that underlying differences in plant physiology or nectar secretion mechanisms across clades more broadly could be responsible for observed differences. Plant clades differ in the location and type of nectaries (Erbar, 2014), resulting in distinct nectar secretion mechanisms and metabolites produced (Göttlinger *et al*., 2024). Mechanisms of nectar production across plant species outside of model organisms are poorly understood (Roy *et al*., 2017) and could underlie the observed correlations among amino acids as well as negative correlations observed between sucrose and amino acids in nectar (Fig 3a). Alternatively, the observed correlations among groups of compounds (Fig 3) may be driven by ecological tradeoffs imposed by pollinators (Adler, 2000; Junker & Blüthgen, 2010), shared biosynthetic pathways (Jenke-Kodama & Dittmann, 2009), or biochemical modification within nectar (Carter & Thornburg, 2004) including the nectar redox cycle, which can lead to generation of pigmentation or other enzymatic or biochemical reactions in nectar.

Many studies have documented variation in microbial abundance in floral nectar across plant species (Herrera *et al*., 2008, 2009b; Alvarez-Perez & Herrera, 2013; Canto *et al*., 2017; Vannette *et al*., 2021; Rering *et al*., 2024). Although variation in microbial dispersal via floral visitors can be an important driver of microbial establishment and growth (Brysch-Herzberg, 2004; Ushio *et al*., 2015; Morris *et al*., 2020), floral chemistry is also linked to variation microbial abundance (Burdon *et al*., 2018; Hanusch *et al*., 2025) and contributions to floral phenotype. Our inoculation approach removed variation imposed by animal-mediated dispersal, by introducing a fixed microbial community across all plant species and examining establishment after 24 hours. Among plant species, nectar amino acid concentration was the strongest predictor of microbial density and diversity in nectar (consistent between individual regressions and module analysis), suggesting that microbial growth of bacteria and yeasts used in this study is nitrogen-limited in floral nectar as previously implicated (Chappell *et al*., 2025). Bacterial use of pollen nitrogen (Christensen *et al*., 2021) is likely one adaptation to this constraint. In contrast to predictions, neither sugar composition, concentration nor pantothenic acid concentration predicted variation in the abundance of the microbial species used here across plant species.

However, we also identified some metabolites linked to reduced microbial growth and diversity. Coexpression analysis suggests that some compounds in nectar enhance microbial diversity, likely by limiting the growth of some otherwise dominant microbial species. Compounds in this module included imidazole alkaloids, fatty aldehydes as well as 2-pyrone derivatives and cyanogenic glycosides (Table S4). Antimicrobial activity has been documented for at least some imidazole alkaloids (Othman *et al*., 2019), while cyanogenic glycosides have diverse functions including modulating oxidative stress (Gleadow & Møller, 2014). In addition to the putative antimicrobial compound classes identified through comparative analyses employed here, we suspect some antimicrobial chemistries may be restricted to plant species or clades and therefore not detectable using our comparative approach (Agrawal & Weber, 2015). For example, *Arbutus unedo* hosted the lowest microbial growth of all species examined here (Cecala *et al*., 2025) and also contains high concentrations of hydrogen peroxide (Landucci & Vannette 2025). Yet this species also uniquely contains the benzenoid bisabolane, a sesquiterpene with demonstrated antibacterial properties (Shu *et al*., 2021). In this case, unique chemistry cannot be definitively linked with antimicrobial activity yet identifying nectars with particularly high or low microbial growth relative to their amino acid concentration may suggest promising bioactive chemistry for further study. Other compounds detected here may contribute to antimicrobial activity in nectar, including widespread coumaric acid or similar phenolics, iridoid glycosides (Whitehead *et al*., 2016) and sesquiterpenoids (Li *et al*., 2022) yet our approach does not point to their strong antimicrobial activity across species. This may be due to their limited effects in nectar, widespread microbial adaptation to such stressors or cogrowth with other microbes. For example, our previous work has shown that the nectar yeast *Metschnikowia reukaufii* is able to grow in nectar containing high levels of peroxides and can detoxify nectar to enable other species to survive (Landucci & Vannette, 2025).

Future work considering the possible contributions and synergies of the measured metabolites with enzymes, ions, volatiles in floral headspace or even physical defenses by plants may lead to increased insights into reward or defensive adaptations associated with particular plant clades or pollination syndromes. Yet it is clear that antimicrobial chemistry of flowers is linked to ecological drivers including floral longevity (Afagwu *et al*., 2026) and can vary among floral tissues and through floral development (Boachon *et al*., 2019). While integrating nectar chemistry with such ecological, physiological and developmental variation is a promising avenue for future research, our study shows that nectar chemical composition can shape the microbial communities within floral nectar and lends further complexity to the chemical signaling and interactions between plants and pollinators mediated by floral nectar.

## Data availability

Raw data and code will be made publicly available on Dryad and Zenodo upon acceptance.

## Acknowledgments

We thank all Vannette lab members including Danielle Rutkowski, Alexia Martin, Shawn Christensen, Dino Sbardellati, Gillian Bergmann, and Jacob Francis for feedback throughout the project. We are grateful to the UC Davis Botanical Conservatory and UC Davis Arboretum staff as well as the UC Davis grounds crew for allowing us to conduct this work with plants on the UC Davis campus. This work is supported in part by an NSF REPS Postbaccalaureate Research Fellowship to LL, a USDA National Institute of Food and Agriculture (NIFA) Postdoctoral Fellowship # 2021-67034-35157 to J.M.C., USDA-ARS Research Project 6036-224340-001 to CR, DEB # 1846266 and Hatch Multistate funding to RLV. The authors declare no conflict of interest.

## Supporting Information

See supporting information for supplementary figures and tables.

