## Supplemental methods for "Comparative nectar metabolomics reveals sucrose-nitrogen tradeoffs and chemical drivers of microbial growth in floral nectar"

*Section S1. Untargeted metabolomics*

Samples were analyzed by liquid chromatography mass spectrometry (LC-MS) in positive and negative mode. A Vanquish LC system equipped with a Hypsersil Gold column (2.1 x 150 mm, 1.9 µm, Thermo Scientific, Waltham, MA, United States) held at 40 °C delivered a linear gradient elution at 0.3 mL min^-1^ as follows: 0% B at 0 min, ramp to 50% B at 5 min, ramp to 98% B at 6 min, hold at 98% B till 10 min, return to 0% B at 10.1 min, total run time: 15 min. A binary mobile phase system was adopted, 0.1% formic acid in A: water and B: acetonitrile. Samples were held at 25 °C in the autosampler and a needle wash with 10% methanol was used before and after each injection (5 s, 50 µL s-1). LC eluent was directed to an IQ-X Orbitrap MS with source conditions as follows: static spray voltage +3.5 kV in positive mode and -2.5 kV in negative mode, gasses (arbitrary units): sheath = 40, auxiliary = 8, sweep = 1, ion transfer tube and vaporizer temperature = 300 °C. Mild trapping mode was enabled, which uses less energy to direct ions through the ion routing multipole, thus limiting MS^1^ fragmentation. Source fragmentation was not enabled. The default charge state was set to 1 and the expected peak width to 6 s. Advanced peak determination algorithm was not used, as most analytes were assumed to be singly charged. The orbitrap full scan ranged from 50-800 m/z and collected spectra in profile mode with resolution set to 120,000 and RF lens at 50%. The quadrupole was used to isolate ions, one microscan was collected per scan, and custom automatic gain control (AGC) mode was used to allow a normalized AGC target of 25%. Auto maximum injection time mode was used, wherein the system calculates and selects the maximum amount of available injection time available given the set scan rate. An internal EASY-IC™ mass calibrant was used to ensure mass accuracy.

We used the AcquireX platform to collect data-dependent MS^2^ spectra. In this process, exclusion and inclusion lists are first generated via MS^1^ analysis of the pooled method blank and pooled sample, respectively. The exclusion list includes *m/z* values not targeted for MS^2^ acquisition while the inclusion list contains features selected for MS^2^ acquisition. Then, iterative MS^2^ analyses of the pooled sample (ID files) are performed using an MS^2^ AcquireX-enabled method. In this process, after precursors on the inclusion list are successfully fragmented, they are automatically moved to the exclusion list and thus, MS^2^ scans are acquired from new and increasingly less abundant features.

A custom AcquireX Deep Scan workflow with legacy component detection algorithm was used. Full scan methods identical to those described previously were created with AcquireX modifications enabled; the MS^2^ method used is reported in the following paragraph. An exclusion override factor was applied such that fragmentation of precursors from the exclusion list was permitted when their abundance was 3-fold greater than the intensity recorded in the pooled blank. No peak window extensions were used for the exclusion or inclusion lists. The inclusion list peak fragmentation threshold was set to 0%. We selected [M+H]^+1^ and [M-H]^-1^ as the preferred ions for fragmentation and used a 6 s exclusion duration, corresponding to predicted peak width. Automatic isotope addition was enabled to exclude isotopes of previously fragmented targets. A 5.0 x 10^4^ intensity threshold and 5 ppm mass tolerance were established.

In our MS^2^ AcquireX-enabled method, we selected data dependent mode = cycle time, wherein as many scans as possible are performed within the specified cycle time, which we set to 1.2 s. MS^1^ orbitrap settings were identical to those used in the full scan except resolution was 60,000. Precursor selection was filtered with an intensity threshold set to 2 x 10^4^ and dynamic mass exclusion applied such that a precursor mass with 3 ppm mass tolerance was excluded for 5 s after fragmentation, and the selected precursor and its isotopes were excluded within a cycle. If no precursors met the target thresholds, the most intense ion was selected for fragmentation. Precursor ions were fragmented by higher-energy collisional dissociation (HCD) and by collision-induced dissociation (CID). For HCD, precursors were delivered using the quadrupole to isolate (1.5 m/z window, no offset) and stepped normalized collision energies (20, 40, 80%) applied. For CID, a 10 ms activation time was used. Product ions from both HCD and CID fragmentation were detected by the orbitrap at 30,000 resolution with auto scan range mode enabled, a standard AGC target and maximum injection time of 54 ms, 1 microscan, and profile mode spectra type. The internal calibrant was not used in MS^2^ scans.

*Section S2. Data processing and feature annotation*

Data were centroided and exported to .mzML format using Compound Discoverer 3.3 (ThermoFisher) for processing using MZmine software version 4.1.0 (Schmid *et al.*, 2023). Spectra were processed by mass detection (MS^1^ noise level filter of 3.0 x 10^4^, no filter applied for MS^2^ mass detection, fragment scans were not denormalized prior to spectral merging). Features were detected using the Chromatogram builder (Myers *et al.*, 2017) with the following parameters: minimum consecutive scans = 4, minimum intensity for consecutive scans = 9 x 10^4^, minimum absolute height = 2.1 x 10^5^, *m/z* tolerance = 0.002 *m/z* or 5 ppm. Chromatograms were smoothed with the Savitzky Golay algorithm. Overlapping features were resolved using the Local minimum resolver algorithm with parameters: MS/MS scan pairing: MS^1^ to MS^2^ precursor tolerance = 0.01 *m/z* or 5 ppm, retention time (RT) filter = 0.5 min, minimum required signals = 1, dimension = RT, chromatographic threshold = 80%, minimum search range RT = 0.041, minimum absolute height = 2.1 x 10^5^, minimum ratio of peak top/edge = 1.80, minimum scans = 4. Isotopes were filtered and grouped with the ^13^C Isotope filtering (intra-sample *m/z* tolerance = 0.001 *m/z* or 5 ppm, RT tolerance = 0.1 min, monotonic shape enabled, maximum charge = 3, representative isotope = most intense, never remove feature with MS^2^ enabled) and isotope peaks finder (chemical elements = H, C, N, O, S, *m/z* tolerance = 0.001 *m/z* or 5 ppm, maximum charge of isotope *m/z* = 3, search in scans = single most intense). Features were aligned across samples with Join aligner (*m/z* tolerance = 0.001 *m/z* or 5 ppm, weight for *m/z* = 3, RT tolerance = 0.2 min, weight for RT = 1). Features were filtered with the Feature list rows filter (minimum aligned features (samples) = 2, minimum features in an isotope pattern = 2, validate ^13^C isotope pattern enabled: *m/z* tolerance = 0.001 *m/z* or 5 ppm, maximum charge = 2, estimate minimum carbon enabled, remove features if ^13^C enabled, exclude isotopes = O, never remove features with MS^2^). Features were grouped with Correlation grouping (i.e., metaCorrelate) according to feature shape (RT tolerance = 0.33 min, intensity threshold for correlation = 3.0 x 10^4^, feature shape correlation enabled with minimum data points = 2, minimum data points on edge = 2, measure = Pearson, minimum feature shape correlation = 85%) and feature height (minimum data points = 2, minimum correlation = 70%). Ion identity molecular networking was used to additionally group features whose precursor ions fulfilled the following: m/z tolerance 0.001 m/z or 5 ppm, check all features, maximum charge = 2, maximum molecules/cluster = 4, positive mode species: [M+H]+, [M+Na]+, [M+K]+, [M+NH_4_]+, [M+2H]2+, [M-H_2_O]+, [M+H-2H_2_O]+, negative mode species: [M-H]-, [M+FA]-, [M-H_2_O]- (Schmid *et al.*, 2021). Ion identity molecular networking annotations were refined by deleting small networks without a major ion and deleting networks with less than 4 links. We used the Export molecular networking files module to generate files (MS^2^ MGF files, feature quantification table with areas, and ion identity edges from IIMN) for feature based molecular networking (FBMN, release 28.2, (Nothias *et al.*, 2020)) via the Global Natural Products Social molecular networking platform (GNPS, (Wang *et al.*, 2016)).

Features retained by MZMine and MS^2^ data were annotated using SIRIUS v6.0.5 (Dührkop *et al.*, 2019) relying on CANOPUS (Dührkop *et al.*, 2021), Classyfire and CSI:FingerID. SIRIUS processing parameters were filtered by isotope pattern with 4 ppm mass accuracy, and de novo and bottom up molecular formula generation were included with elements H, C, N, O, and P included. Processing settings for other annotation features followed default settings. CANOPUS is a deep neural network that classifiers unknown metabolites based on MS^2^ fragmentation patterns. Annotations were combined into a feature table for downstream analyses. Untargeted compound diversity within a sample was estimated using Hill number N=1 (Chao *et al.*, 2014).

*Section S3. Targeted metabolomics – Carbohydrates*

Nectar carbohydrates were analyzed via targeted analysis with a Vanquish LC system coupled to a TSQ Quantis™ MS (Thermo Scientific). Sugars were separated on a Hypersil GOLD™ Amino column (100 x 2.1 mm, 1.9 μm, Thermo Scientific) held at 25 °C and isocratic mobile phase delivered at 0.5 ml min^-1^ of 2 mM ammonium acetate in 80:20 acetonitrile/water adjusted to pH = 5.4 with acetic acid. LC effluent was vaporized via electrospray ionization in negative mode at 4500 V. The vaporizer temperature was held at 400 °C and sheath, auxiliary, and sweep gases set to 60, 15, and 2 arbitrary units, respectively. Single reaction monitoring mode was used to detect sugars as reported in Table S1. A cycle time of 0.8 seconds was used with a chromatographic peak width of 10 s. Collision induced dissociation gas (argon) was provided at 1.5 mTorr and Q1 and Q3 resolution were 0.7 and 1.2 full width at half maximum, respectively. For major sugars (glucose, fructose, and sucrose), a 3 min run time was used and samples were injected after 2500-fold dilution. Minor sugar components had a run time of 16 min and 250-fold dilution. External calibration standards were used to quantify sugars according to their quantitative transition’s peak area. Calibration standard ranges were selected to encompass the abundance of sample peak areas.

**Table S1. Carbohydrate analytical details.** Minor and major carbohydrate components of nectar are listed with their retention times and precursor (representing [M-H]^-^) and product ion pairs. Product ions selected as quantitative transitions score peak areas in samples and calibration standards are indicated in bold with an asterisk. Collision energies for each respective transition are provided. External calibration standard concentration ranges and the resulting calibration curves’ coefficients of determination (R^2^) are provided. The lowest calibration standard represents the limit of quantification in our analysis.

| **Group** | **Compound** | **Retention time (min)** | **Precursor ion ([M-H]^-^; *m/z*)** | **Product ion (*m/z*)** | **Collision energy (V)** | **RF lens (V)** | **Calibration curve range** | **R^2^** |
| --- | --- | --- | --- | --- | --- | --- | --- | --- |
| Minor carbohydrate components | Myo-inositol | 2.08 | -179.0 | **-161.0*** | 10.35 | 87 | 1 - 50 mg/L | 0.9979 |
|  |  |  |  | -87.0 | 15.28 |  |  |  |
|  |  |  |  | -98.9 | 16.08 |  |  |  |
|  |  |  |  | -117.0 | 12.33 |  |  |  |
|  | Erythritol | 0.97 | -121.0 | **-89.0*** | 8.79 | 53 | 1 - 25 mg/L | 0.9996 |
|  |  |  |  | -71.0 | 13.72 |  |  |  |
|  |  |  |  | -101.0 | 6.47 |  |  |  |
|  | Maltose | 2.1 | -341.0 | **-161.0*** | 5.25 | 75 | 1 - 50 mg/L | 0.9979 |
|  |  |  |  | -101.0 | 13.3 |  |  |  |
|  |  |  |  | -179.0 | 5.46 |  |  |  |
|  |  |  |  | -221.0 | 15.15 |  |  |  |
|  | Maltotriose | 3.58 | -503.2 | **-341.1*** | 6.14 | 124 | 1 - 25 mg/L | 0.9987 |
|  |  |  |  | -179.0 | 10.73 |  |  |  |
|  | Mannitol/sorbitol | 1.36 | -181.0 | **-89.1*** | 13.09 | 77 | 1 - 100 mg/L | 0.9999 |
|  |  |  |  | -59.0 | 18.82 |  |  |  |
|  |  |  |  | -100.9 | 12.96 |  |  |  |
|  | Melezitose | 3.03 | -503.1 | **-323.1*** | 18.06 | 181 | 1 - 50 mg/L | 0.9994 |
|  |  |  |  | -179.0 | 18.94 |  |  |  |
|  | Melibiose | 2.72 | -341.0 | **-179.0*** | 7.7 | 77 | 1 - 25 mg/L | 0.9969 |
|  |  |  |  | -221.0 | 10.18 |  |  |  |
|  |  |  |  | -251.1 | 5.76 |  |  |  |
|  | Raffinose | 3.68 | -503.1 | **-179.0*** | 19.74 | 178 | 1 - 5 mg/L | 0.9746 |
|  |  |  |  | -221.1 | 26.27 |  |  |  |
|  |  |  |  | -323.0 | 17.34 |  |  |  |
|  | Ribose | 0.8 | -149.0 | **-89.0*** | 5.25 | 53 | 1 - 50 mg/L | 0.9999 |
|  |  |  |  | -59.0 | 12.62 |  |  |  |
|  |  |  |  | -71.0 | 8.62 |  |  |  |
|  | Stachyose | 8 | -665.2 | **-383.1*** | 31.03 | 231 | 1 - 50 mg/L | 0.9999 |
|  |  |  |  | -341.1 | 24.42 |  |  |  |
|  |  |  |  | -485.1 | 25.26 |  |  |  |
|  | Trehalose | 2.3 | -401.1 | **-341.1*** | 12.67 | 142 | 1 - 5 mg/L | 0.9664 |
|  |  |  |  | -119.1 | 25.22 |  |  |  |
|  |  |  |  | -161.1 | 21.51 |  |  |  |
|  |  |  |  | -179.1 | 19.87 |  |  |  |
|  | Xylitol | 1.16 | -151.0 | **-89.0*** | 11.07 | 69 | 1 - 25 mg/L | 0.9764 |
|  |  |  |  | -59.0 | 16.58 |  |  |  |
| Major carbohydrate components | Fructose | 1.1 | -179.0 | **-89.0*** | 7.57 | 87 | 1 - 100 mg/L | 0.9209 |
|  |  |  |  | -59.0 | 16.54 |  |  |  |
|  |  |  |  | -71.0 | 14.98 |  |  |  |
|  | Glucose | 1.22 | -179.0 | **-89.0*** | 6.22 | 30 | 1 - 100 mg/L | 0.9982 |
|  |  |  |  | -59.1 | 16.33 |  |  |  |
|  |  |  |  | -119.0 | 5.25 |  |  |  |
|  | Sucrose | 1.59 | -341.1 | **-179.0*** | 13.47 | 125 | 1 - 100 mg/L | 0.9953 |
|  |  |  |  | -89.1 | 20.63 |  |  |  |
|  |  |  |  | -113.1 | 20.16 |  |  |  |
|  |  |  |  | -119.0 | 18.48 |  |  |  |

**Table S2. Analytical details for amino acid and pantothenic acid post hoc quantification.**

| **Compound** | **Retention time** | **Quantitative ion; [M+H]^+^** | **Calibration curve range** | **R^2^** |
| --- | --- | --- | --- | --- |
|  | **(min)** | **(*m/z*)** | **(µg/L)** |  |
| Alanine | 1.35 | 90.055 | 5-500 | 0.9999 |
| Arginine | 1.29 | 175.119 | 5-500 | 0.9961 |
| Asparagine | 1.31 | 133.0608 | 1-500 | >0.9999 |
| Aspartic acid | 1.32 | 134.0448 | 5-500 | 0.9999 |
| Glutamic acid | 1.37 | 148.0604 | 1-500 | >0.9999 |
| Glutamine | 1.34 | 147.0764 | 1-500 | 0.9988 |
| Histidine | 1.25 | 156.0768 | 5-500 | 0.9994 |
| Hydroxyproline | 1.36 | 132.0655 | 5-500 | 0.9998 |
| Isoleucine | 2.88 | 132.1019 | 5-500 | 0.9999 |
| Leucine | 2.95 | 132.1019 | 5-500 | 0.9997 |
| Lysine | 1.22 | 147.1128 | 5-500 | 0.9971 |
| Methionine | 2.31 | 150.0583 | 5-500 | 0.9999 |
| Ornithine | 1.22 | 133.0972 | 10-500 | 0.9963 |
| Pantothenic acid | 3.37 | 220.1175 | 25-500 | >0.9999 |
| Phenylalanine | 3.31 | 166.0863 | 1-500 | >0.9999 |
| Proline | 1.56 | 116.0706 | 5-500 | >0.9999 |
| Threonine/ homoserine* | 1.34 | 120.0655 | 1-500 | >0.9999 |
| Tryptophan | 3.79 | 205.0972 | 1-500 | 0.9999 |
| Tyrosine | 2.78 | 182.0812 | 5-500 | >0.9999 |
| α-Aminobutyric acid (AABA) | 1.56 | 104.0706 | 10-500 | 0.9984 |
| γ-Aminobutyric acid (GABA) | 1.41 | 104.0706 | 5-500 | 0.9998 |

* We were able to detect 21 amino acids including threonine and homoserine although these could not be resolved as individual features so their abundances are combined.
