## Supplementary Figures for "Comparative nectar metabolomics reveals sucrose-nitrogen tradeoffs and chemical drivers of microbial growth in floral nectar"

Supplementary figures and tables for “Comparative nectar metabolomics reveals sucrose-nitrogen tradeoffs and chemical drivers of microbial growth in floral nectar”

**Table S1**. Classification of annotated compounds detected in positive mode using Classyfire as implemented in CANOPUS and SIRIUS. Feature richness is across all samples in the dataset after blank subtraction.

| **Classyfire superclass** | **Richness** | **Class** |
| --- | --- | --- |
| Alkaloids and derivatives | 3 | Alkaloids and derivatives, Aporphines |
| Benzenoids | 26 | 2(hydroxyphenyl)acetic acids, Benzenoids, Benzene and substituted derivatives, Benzoic acids and derivatives, Anisoles, Benzoyl derivatives, Galloyl esters, Acylsalicylic acids, Methoxyphenols, Naphthalenes |
| Hydrocarbons | 2 | Acetylenes |
| Lipids and lipid-like molecules | 71 | Fatty acyl glycosides of mono- and disaccharides, Nitro fatty acids, Unsaturated fatty acids, Fatty acid esters, Alkyl glycosides, Thia fatty acids, Iridoid O-glycosides, Acyl carnitines, Terpene glycosides, Fatty Acyls, Abscisic acids and derivatives, Long-chain fatty acids, N-acyl amines, Fatty alcohols, Fatty amides, Sesquiterpenoids, Phosphosphingolipids, Lipids and lipid-like molecules, Fatty acids and conjugates, Phosphatidylcholines, Dialkylglycerols, Ceramides |
| Nucleosides, nucleotides, and analogues | 1 | 1-ribosyl-imidazolecarboxamides |
| Organic 1,3-dipolar compounds | 2 | Organic nitro compounds |
| Organic acids and derivatives | 98 | Alpha amino acids, L-alpha-amino acids, Histidine and derivatives, Oligopeptides, Alpha amino acids and derivatives, D-alpha-amino acids, Amino acids, Carboxylic acid derivatives, Monocarboxylic acids and derivatives, Carboxylic acid esters, Dicarboxylic acids and derivatives, Carboxylic acids, Carboxylic acids and derivatives, Peptides, Amino acids and derivatives, Hydroxy acids and derivatives, Short-chain hydroxy acids and derivatives, Alpha hydroxy acids and derivatives, Proline and derivatives, Alpha amino acid esters, Phenylalanine and derivatives, Kainoids, Beta amino acids and derivatives, N-acyl-L-alpha-amino acids, Monoalkyl phosphates, N-acyl-alpha amino acids and derivatives, N-acyl-alpha amino acids, Thiocarboxylic acids and derivatives, Arylsulfates, Carboxylic acid amides, Tertiary carboxylic acid amides |
| Organic nitrogen compounds | 21 | Guanidines, Phosphocholines, Tetraalkylammonium salts, Dialkylamines, Amines, Monoalkylamines, 1,2-aminoalcohols |
| Organic oxygen compounds | 74 | Primary alcohols, Monosaccharides, Hexoses, Enals, Carbonyl compounds, C-glycosyl compounds, Oligosaccharides, Disaccharides, Acylaminosugars, Acyloins, Cyclohexanols, Glycosylamines, Alpha,beta-unsaturated aldehydes, Cyanogenic glycosides, Glycosyl compounds, Glucuronic acid derivatives, Polyethylene glycols, Phenolic glycosides, Secondary alcohols, Ketones, Alkyl aryl ethers |
| Organoheterocyclic compounds | 38 | Azoles, Pyranones and derivatives, Oxacyclic compounds, Butenolides, Pyridinecarboxylic acids, Gamma butyrolactones, Lactones, Furans, Diazines, Lactams, Pyrimidones, Indolecarboxylic acids and derivatives, Benzoquinolines, Indoles, Aminopyrimidines and derivatives, Azacyclic compounds, Thiepanes, Pyrimidines and pyrimidine derivatives, Heteroaromatic compounds, Aminotriazines |
| Organophosphorus compounds | 2 | Organophosphorus compounds |
| Organosulfur compounds | 3 | Thioethers, Sulfenyl compounds, Dialkylthioethers |
| Phenylpropanoids and polyketides | 11 | Coumaric acids, Phenylpropanoic acids, Flavonoid O-glycosides, Cinnamaldehydes, Coumaric acids and derivatives, Flavonols |

**Table S2.** Classification of annotated compounds detected in negative mode using NP Classifier as implemented in CANOPUS and SIRIUS. Feature richness is across all samples in the dataset after blank subtraction.

| **NPC.pathway..** | **Richness** | **Class** |
| --- | --- | --- |
| Alkaloids | 29 | Imidazole alkaloids, Purine alkaloids, Purine nucleosides, Pyridine alkaloids, Cyanogenic glycosides, Acridone alkaloids, Amino acids, Polyamines, Cytochalasan alkaloids |
| Amino acids and Peptides | 55 | Amino acids, Imidazole alkaloids, Glucosinolates, Cyanogenic glycosides, Dipeptides, Leukotrienes, Cephalosporins, Melithiazole and Myxothiazole derivatives, Cephamycins, Penicillins |
| Carbohydrates | 57 | Monosaccharides, Cyclitols, Aminosugars, Disaccharides, Polysaccharides, Purine nucleosides, Pyrimidine nucleosides, Purine nucleotides, Imidazole alkaloids |
| Fatty acids | 56 | Halogenated fatty acids, Thia fatty acids, Dicarboxylic acids, Fatty alcohols, Monosaccharides, Unsaturated fatty acids, Oxo fatty acids, Fatty aldehydes, Hydroxy fatty acids, Wax monoesters, N-acyl amines, Branched fatty acids, Fatty acyl glycosides of mono- and disaccharides, Other Octadecanoids, Amino acids, Methoxy fatty acids |
| Polyketides | 4 | Naphthoquinones, Tropolones and derivatives (PKS), Anthraquinones and anthrones, Open-chain polyketides |
| Shikimates and Phenylpropanoids | 38 | Simple phenolic acids, Isoflavones, Shikimic acids and derivatives, Gallotannins, Phenylethanoids, Flavan-3-ols, Cinnamic acids and derivatives, Phthalide derivatives, Flavones, Flavonols, Dihydroflavonols, Chalcones, Anthraquinones and anthrones |
| Terpenoids | 22 | Iridoids monoterpenoids, Secoiridoid monoterpenoids, Apocarotenoids(ε-), Camphane monoterpenoids, Herbertane sesquiterpenoids |

**Table S3.** Chemical compounds recovered in negative mode using Classyfire as implemented in CANOPUS and SIRIUS. Feature richness is across all samples in the dataset after blank subtraction.

| **ClassyFire.superclass..** | **Richness** | **Class** |
| --- | --- | --- |
| Benzenoids | 18 | Benzenoids, Galloyl esters, 1-hydroxy-2-unsubstituted benzenoids, Anthracenecarboxylic acids, Gallic acid and derivatives, m-Hydroxybenzoic acid esters, Salicylic acids, Anthraquinones, Benzamides, Benzenesulfonamides |
| Lipids and lipid-like molecules | 40 | Straight chain fatty acids, Fatty acyl glycosides of mono- and disaccharides, Fatty acyl glycosides, Fatty Acyls, Alkyl glycosides, Iridoid O-glycosides, Methyl-branched fatty acids, Terpene glycosides, Jasmonic acids, Fatty alcohols, Long-chain fatty acids, Bicyclic monoterpenoids, Sesquiterpenoids, Linoleic acids and derivatives, Medium-chain fatty acids |
| Nucleosides, nucleotides, and analogues | 7 | Triazole ribonucleosides and ribonucleotides, Pyrimidine nucleosides, Purine nucleosides, Pyrimidine ribonucleotides |
| Organic 1,3-dipolar compounds | 1 | Nitroaromatic compounds |
| Organic acids and derivatives | 79 | Alpha amino acids and derivatives, Organic phosphoric acids and derivatives, Carboxylic acids, Alkyl phosphates, Histidine and derivatives, Carboxylic acid derivatives, Beta hydroxy acids and derivatives, N-acyl ureas, Alpha amino acids, L-alpha-amino acids, Medium-chain hydroxy acids and derivatives, Alpha-keto acids and derivatives, Alpha hydroxy acids and derivatives, Dicarboxylic acids and derivatives, N-carbamoyl-alpha amino acids, Medium-chain keto acids and derivatives, Organosulfonic acids and derivatives, Tricarboxylic acids and derivatives, N-acyl-alpha amino acids and derivatives, N-acyl-alpha amino acids, Beta amino acids and derivatives, Phenylsulfates, Organosulfonamides, S-alkyl thiosulfates, Sulfuric acid monoesters |
| Organic nitrogen compounds | 3 | N-arylamides |
| Organic oxygen compounds | 55 | Secondary alcohols, Inositol phosphates, Hexoses, Monosaccharides, Monosaccharide phosphates, O-glycosyl compounds, C-glucuronides, Glucuronic acid derivatives, Ketones, C-glycosyl compounds, O-glucuronides, Disaccharides, Disaccharide sulfates, Glucosinolates, Oligosaccharides, Cyclic ketones, Beta-hydroxy aldehydes, Glucuronides, Quinic acids and derivatives, Phenolic glycosides, Glycosylamines, Hydroxybenzaldehydes, Cyclohexenones, Alkyl aryl ethers |
| Organoheterocyclic compounds | 31 | Heteroaromatic compounds, Azacyclic compounds, Azoles, Benzopyrans, Pyridines and derivatives, Nitroquinolines and derivatives, Quinoline carboxylic acids, Purines and purine derivatives, 1-benzopyrans, Aminotriazines, Quinolines and derivatives, Furoic acid and derivatives, 1-benzothiopyrans, N-aliphatic s-triazines, Isoindolones |
| Organosulfur compounds | 7 | Alkylthiols, Sulfenyl compounds, Alkylarylthioethers, Sulfonyls |
| Phenylpropanoids and polyketides | 20 | Isoflavones, Catechins, Hydrolyzable tannins, Hydroxycinnamic acid glycosides, Flavonoid 8-C-glycosides, Coumaric acid esters, Phenylpropanoic acids, Flavonoid C-glycosides, Flavonoid-3-O-glycosides, 3-sulfated flavonoids, Coumaric acids, 3'-hydroxyflavonoids, 5-hydroxyflavonoids |

**Table S4.** Compounds from the untargeted metabolite dataset included in modules that had significant associations with microbial growth in nectar or nectar volume (See Figure S10). Specific names may be missing for some compounds although all are annotated at the level of NPC class.

| **Module** | **ID** | **NPC.class** | | **name** |
| --- | --- | --- | --- | --- |
| black | 1557 | Amino acids | | Harg (arginine) |
| black | 1616 | Amino acids | | (2S)-3-(1H-imidazol-5-yl)-2-(methylamino)propanoate |
| black | 1624 | Amino acids | | Serine |
| black | 1916 | Amino acids | | Hgln (glutamine) |
| black | 1957 | Amino acids | | L-Glu (glutamic acid) |
| black | 2038 | Amino acids | | L-Thr (threonine) |
| black | 2079 | Amino acids | | 2-aminobut-3-enoic acid |
| black | 2135 | Amino acids | | L-Glu (glutamic acid) |
| black | 3771 | Acetate-derived alkaloids | | Cyclopropanemethylamine |
| black | 3779 | Aminoacids | | Hval (valine) |
| black | 5104 | Hydrocarbons | | Cyclopropylmethylium |
| black | 5105 | Aminoacids | | Hval (valine) |
| black | 6166 | Fatty alcohols | | 1-(butylphosphonoyl)butane |
| black | 7630 | Hydrocarbons | | 2-Pentyne |
| black | 7638 | Aminoacids | | Alle (alloisoluecine) |
| black | 7960 | Aminoacids | | Alle (alloisoluecine) |
| black | 7980 | Acetate-derived alkaloids | | (1-Methylcyclobutyl)azanium |
| black | 8572 | Pyrrolidine alkaloids | | (1S,2S,4R,5R,6S)-2-Amino-4-methyl-bicyclo[3.1.0]hexane-2,6-dicarboxylic acid |
| black | 9144 | N-acyl amines | | Pantothenate (pantothenic acid) |
| black | 27717 | Aminoglycosides | | 3-[2-[2-[2-[2-[[2-[(2R)-2-amino-3-oxopropyl]sulfanylacetyl]amino]ethoxy]ethoxy]ethoxy]ethoxy]propanoate |
| Yellow | 1033 | Amino acids | | Lysine |
| Yellow | 1088 | Amino acids | | 2-aminobut-3-enoic acid |
| Yellow | 1125 | Amino acids | | 2-amino-4-(ethylsulfanyl)butanoic acid |
| Yellow | 1375 | Amino acids | | L-His (histidine) |
| Yellow | 1735 | Amino acids | | (2S)-4-oxoazetidine-2-carboxylic acid |
| Yellow | 1782 | Amino acids | | 3-Nitroso-2-oxazolidinone |
| Yellow | 2478 | Amino acids | | (carboxymethyl)trimethylazanium |
| Yellow | 2869 | Pyridine alkaloids | | 2-Methylpyridine-3-carboxylate |
| Yellow | 4948 | Amino acids | | NA |
| Yellow | 5769 | Thia fatty acids | | Thiolane-3-carboxylic acid |
| Yellow | 5770 | Thia fatty acids | | Thiolane-3-carboxylic acid |
| Yellow | 6133 | Amino acids | | NA |
| Yellow | 7180 | Amino acids | | Pidolic Acid (L-Pyroglutamic acid) |
| Yellow | 9253 | Carboline alkaloids | | methyl-1-(2,2-dimethylindan-1S-yl)-imidazole-5-carboxylate |
| Yellow | 11009 | Isoquinoline alkaloids | | 6-O-Methylhaemanthidine |
| Yellow | 11056 | Simple indole alkaloids | | Indoleacrylic Acid |
| Yellow | 11065 | Simple indole alkaloids | | (2R)-3-(1H-indol-3-yl)-2-(trimethylazaniumyl)propanoate (hypaphorine) |
| Yellow | 11066 | Simple indole alkaloids | | (2R)-3-(1H-indol-3-yl)-2-(trimethylazaniumyl)propanoate (hypaphorine) |
| Yellow | 11069 | Purine alkaloids | | 3-azanyl-5-(azepan-1-yl)-N-carbamimidoyl-6-(2-methoxypyrimidin-5-yl)pyrazine-2-carboxamide |
| Yellow | 11070 | Simple indole alkaloids | | (2R)-3-(1H-indol-3-yl)-2-(trimethylazaniumyl)propanoate (hypaphorine) |
| Yellow | 12706 | Isoquinoline alkaloids | | Norboldine |
| Yellow | 12972 | Isoquinoline alkaloids | | NA |
| Yellow | 12974 | Quinoline alkaloids | | benzyl 2-{[(benzyloxy)carbonyl]amino}-3-hydroxypropanoate |
| Yellow | 13569 | Isoquinoline alkaloids | | Erythraline |
| Yellow | 13612 | Isoquinoline alkaloids | | Norboldine |
| Purple | 2368 | Disaccharides | | Umbelliferose |
| Purple | 2836 | Disaccharides | | Sucrose |
| Purple | 3048 | Dicarboxylic acids | | Aminooxy(triaminomethyl)phosphinic acid |
| Purple | 4607 | Disaccharides | | Raffinose |
| Purple | 4608 | Monosaccharides | | NA |
| Purple | 4609 | Cyanogenic glycosides | | [(3aR,6aR)-3-(methylcarbamoyloxy)-2,3,3a,5,6,6a-hexahydrofuro[3,2-b]furan-6-yl] N-[[[(3aR,6aR)-3-(methylcarbamoyloxy)-2,3,3a,5,6,6a-hexahydrofuro[3,2-b]furan-6-yl]oxycarbonylamino]methyl]carbamate |
| Purple | 4612 | Disaccharides | | Lactosucrose |
| Purple | 5359 | Disaccharides | | Lactosucrose |
| Purple | 6832 | Disaccharides | | (2R,3R,4S,5R,6S)-3,4,5-trihydroxy-2-(hydroxymethyl)-6-[(2R,3S,4R,5R,6R)-4,5,6-trihydroxy-2-(hydroxymethyl)oxan-3-yl]oxyoxane-2,3-dicarboxylic acid |
| Purple | 20290 | Primary amides | NA | |
| Purple | 25264 | Fatty alcohols | | Triethylene glycol monotridecyl ether |
| Purple | 29981 | Purine alkaloids | | N-[3-(2-amino-1H-imidazol-4-yl)prop-2-en-1-yl]-1H-pyrrole-2-carboxamide |
| Green | 1869 | Polysaccharides | | NA |
| Green | 1875 | Imidazole alkaloids | | 3-amino-N-ethyl-1,2,4-triazole-1-sulfonamide |
| Green | 2938 | Disaccharides | | NA |
| Green | 3002 | Disaccharides | | 2-hydroxy-2-(4-hydroxy-3-{[3,4,5-trihydroxy-6-(hydroxymethyl)oxan-2-yl]oxy}oxolan-2-yl)acetaldehyde |
| Green | 3017 | Fatty aldehydes | | Butenolide |
| Green | 3076 | 2-pyrone derivatives | | 5-Hydroxy-4-Methoxy-5,6-Dihydro-2H-Pyran-2-One |
| Green | 3089 | Disaccharides | | 2-hydroxy-2-(4-hydroxy-3-{[3,4,5-trihydroxy-6-(hydroxymethyl)oxan-2-yl]oxy}oxolan-2-yl)acetaldehyde |
| Green | 3098 | Cyanogenic glycosides | | 6-(2-Hydroxyethylamino)-3-[3,4,5-trihydroxy-6-(hydroxymethyl)oxan-2-yl]oxyhexane-1,2,4,5-tetrol |
| Green | 3133 | 2-pyrone derivatives | | 4-methoxy-2,5-dihydrofuran-2-one |
| Green | 3137 | Disaccharides | | Brachiose (isomaltose) |
| Green | 3167 | Disaccharides | | NA |
| Green | 3166 | Dicarboxylic acids | | 2,3-Methylenesuccinic Acid |
| Green | 3176 | 2-pyrone derivatives | | (2z)-2-(5-oxofuran-2-ylidene)acetate (cis-dienelactone) |
| Green | 3182 | Disaccharides | | 2,3,5,6-tetrahydroxy-4-{[3,4,5-trihydroxy-6-(hydroxymethyl)oxan-2-yl]oxy}hexanal (maltose hydrate) |
| Green | 3192 | Disaccharides | | NA |
| Green | 3193 | Disaccharides | | Brachiose (isomaltose) |
| Green | 3222 | Disaccharides | | Sucrose |
| Green | 3233 | Purine nucleosides | | 3-[4-(methylsulfanyl)-2-(3-{3,5,9-trimethyl-7-oxo-7H-furo[3,2-g]chromen-6-yl}propanamido)butanamido]propanoic acid |
| Green | 3531 | Purine alkaloids | | NA |
| Green | 6178 | Purine nucleosides | | NA |
| Green | 6976 | Simple phenolic acids | | Ethylylidene triacetate |
| Green | 7043 | Disaccharides | | [(3S,4S,6R)-3,4,5-trihydroxy-6-[(2R,4R)-3,4,5-trihydroxy-6-(hydroxymethyl)oxan-2-yl]oxyoxan-2-yl]methyl acetate |
| Green | 10452 | Pyrrolizidine alkaloids | | 7-({[2,3-dihydroxy-2-(propan-2-yl)butanoyl]oxy}methyl)-1-hydroxy-1,2,3,4,5,7a-hexahydropyrrolizin-4-ium-4-olate |

**Supplementary Figures**

**Figure S1.** Molecular network of features detected in the nectar of 31 phylogenetically diverse plant species. Features are color coded by NP pathway chemical classification annotation and network edges represent structural similarity among compounds based on GNPS.

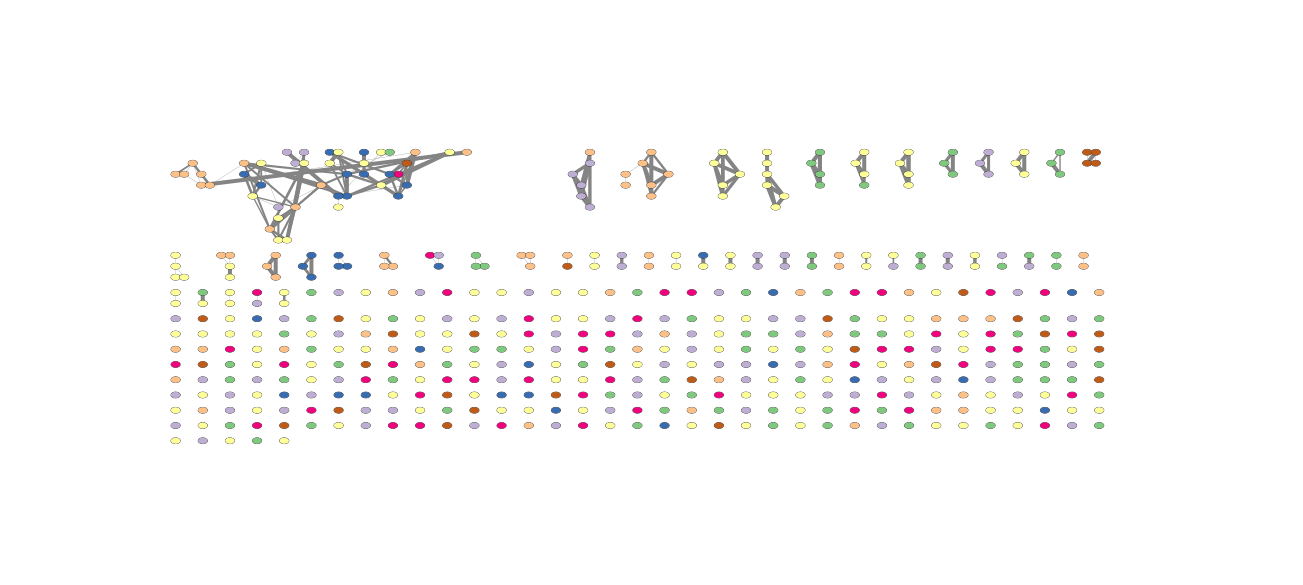

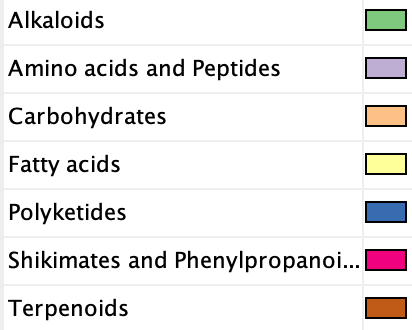

**Figure S2.** Plant clades differ in targeted metabolites including sugars and amino acids.

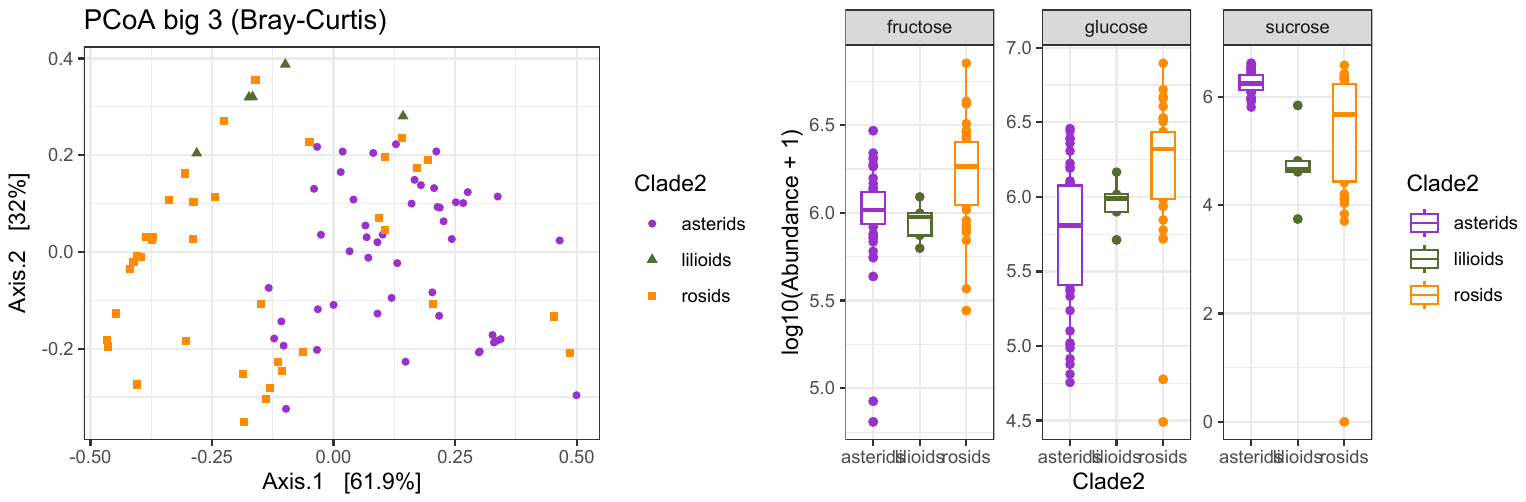

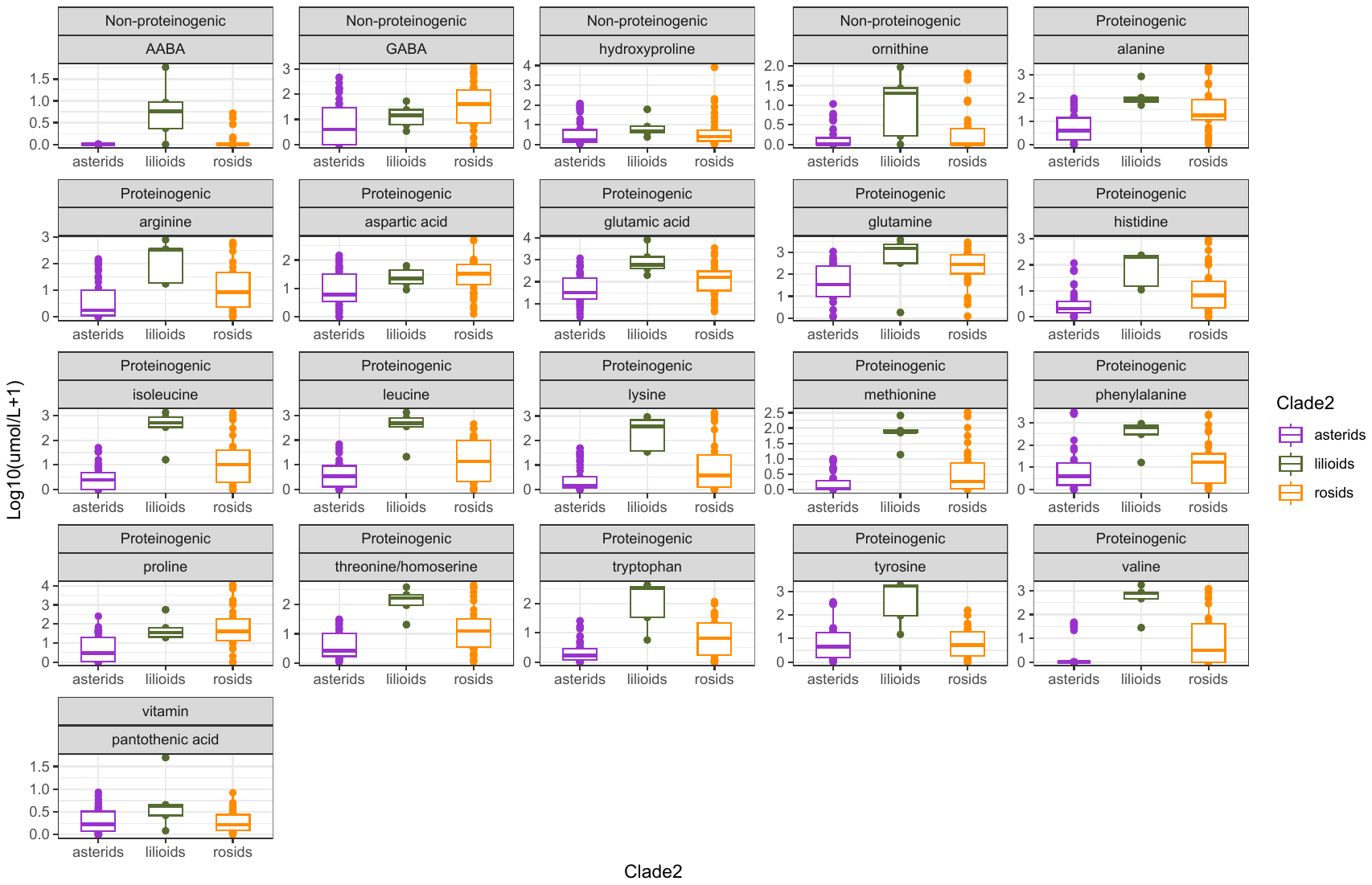

**Figure S3.** Plant clades differ in the concentration of minor sugars and sugar alcohols.
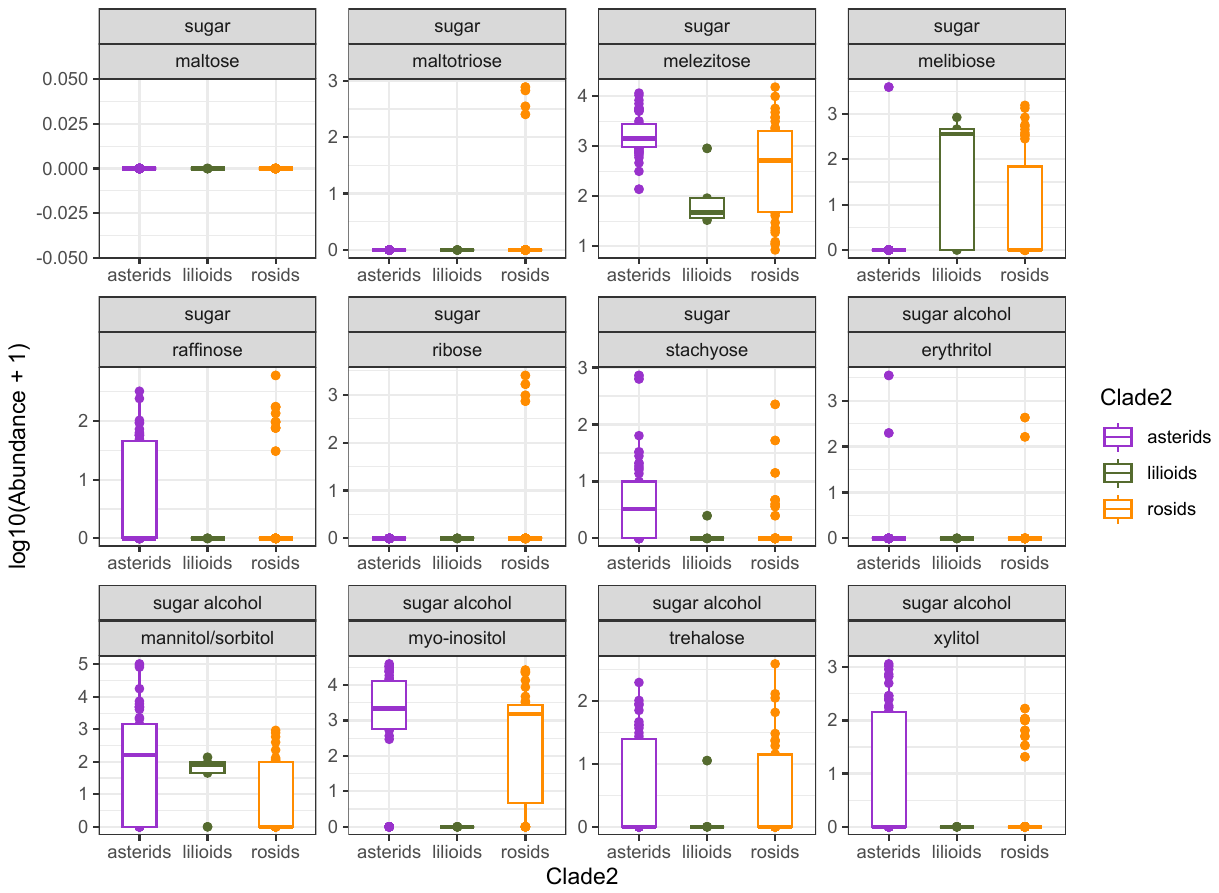

**Figure S4.** Variation among plant species in the composition of Alkaloids (untargeted metabolomics, positive mode). Red indicates high relative abundance, blue indicates not detected.

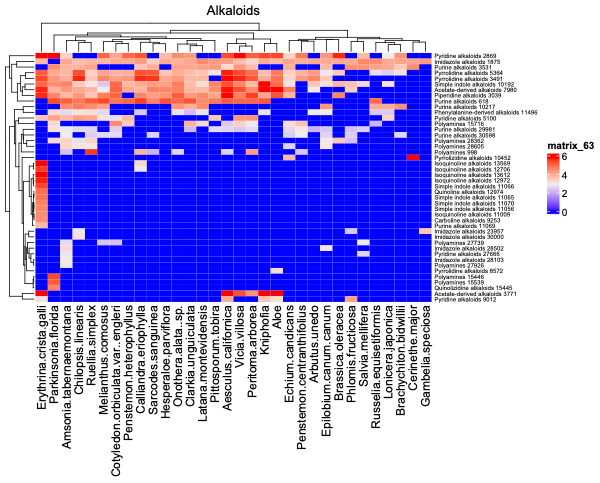

**Figure S5.** Variation among plant species in the composition of Carbohydrates (untargeted metabolomics). Red indicates high relative abundance, blue indicates not detected.

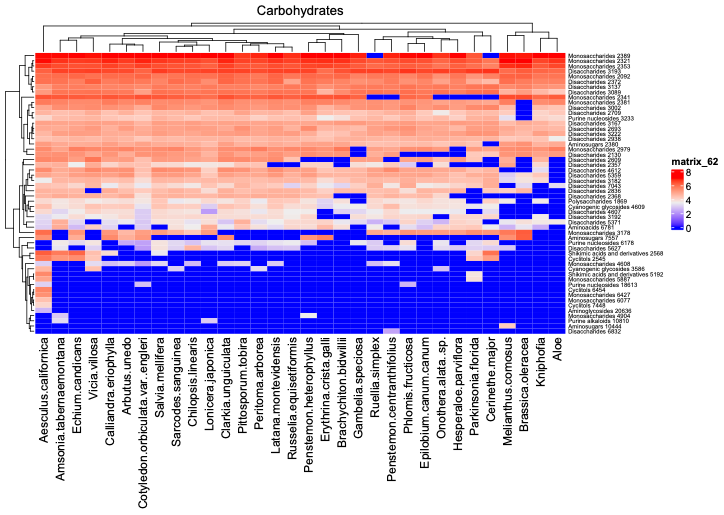

**Figure S6.** Variation among plant species in the composition of Fatty Acids (untargeted metabolomics). Red indicates high relative abundance, blue indicates not detected.

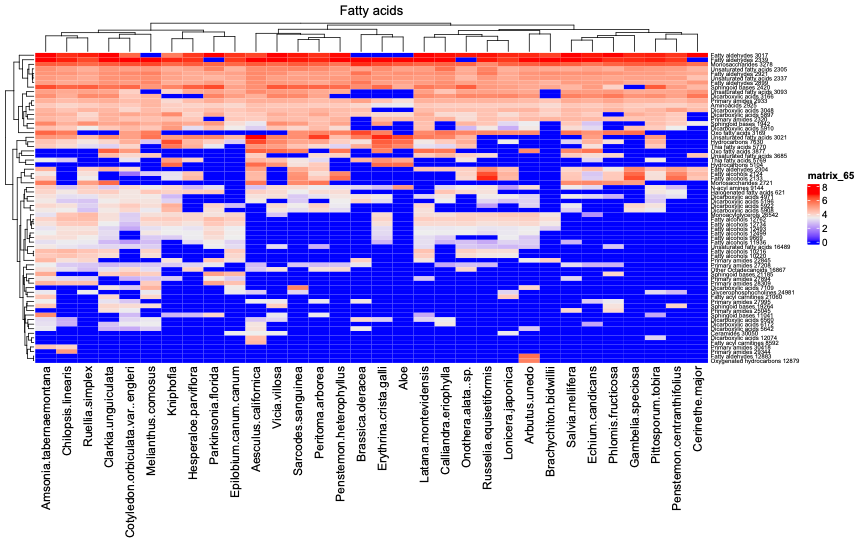

**Figure S7.** Variation among plant species in the composition of Terpenoids (untargeted metabolomics). Red indicates high relative abundance, blue indicates not detected.

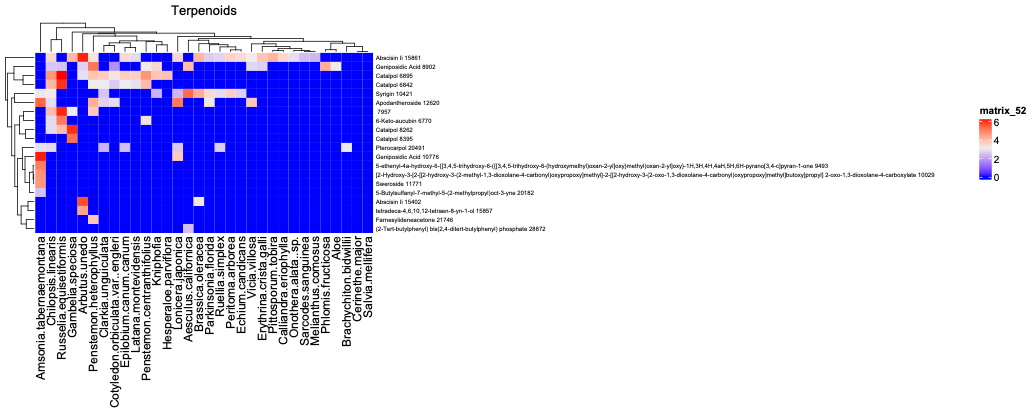

**Figure S8.** Variation among plant species in the composition of Shikimates and Phenylpropanoids (untargeted metabolomics). Red indicates high relative abundance, blue indicates not detected.

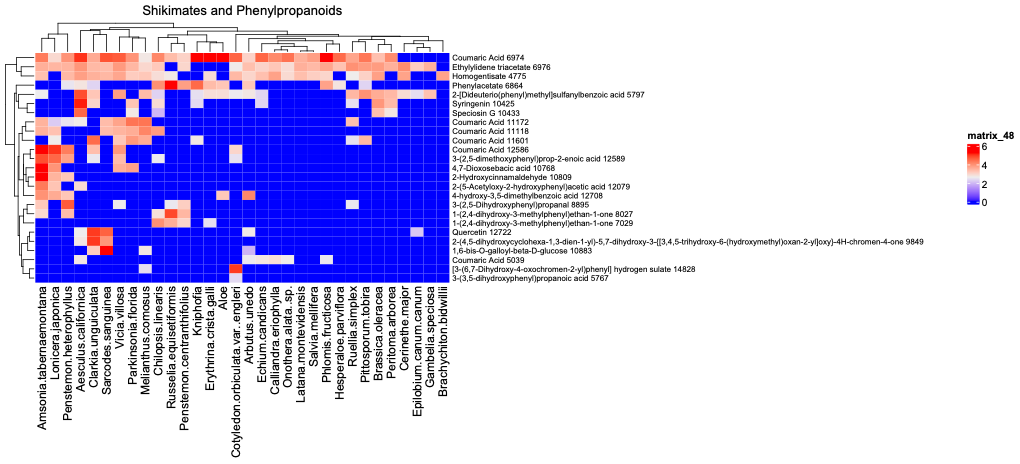

**Figure S9.** Compound dendrogram associated with modules used to predict variation in microbial growth.

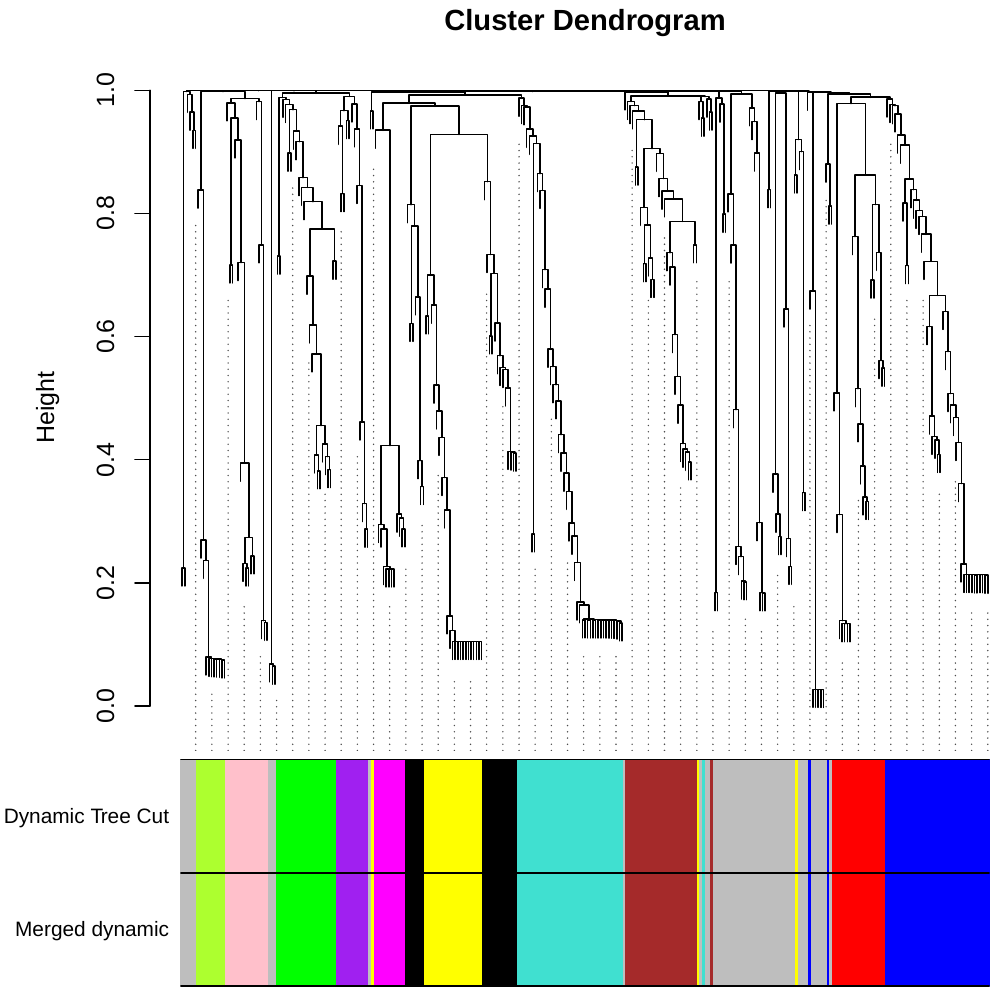

**Figure S10.** Metabolite composition is associated with variation in nectar traits and microbial growth in nectar across plant species. Correlations between metabolite clusters (colors) and measured plant traits, including nectar volume, microbial Shannon diversity within floral nectar (24 hours post inoculation) and microbial density within floral nectar (24 hours post inoculation).

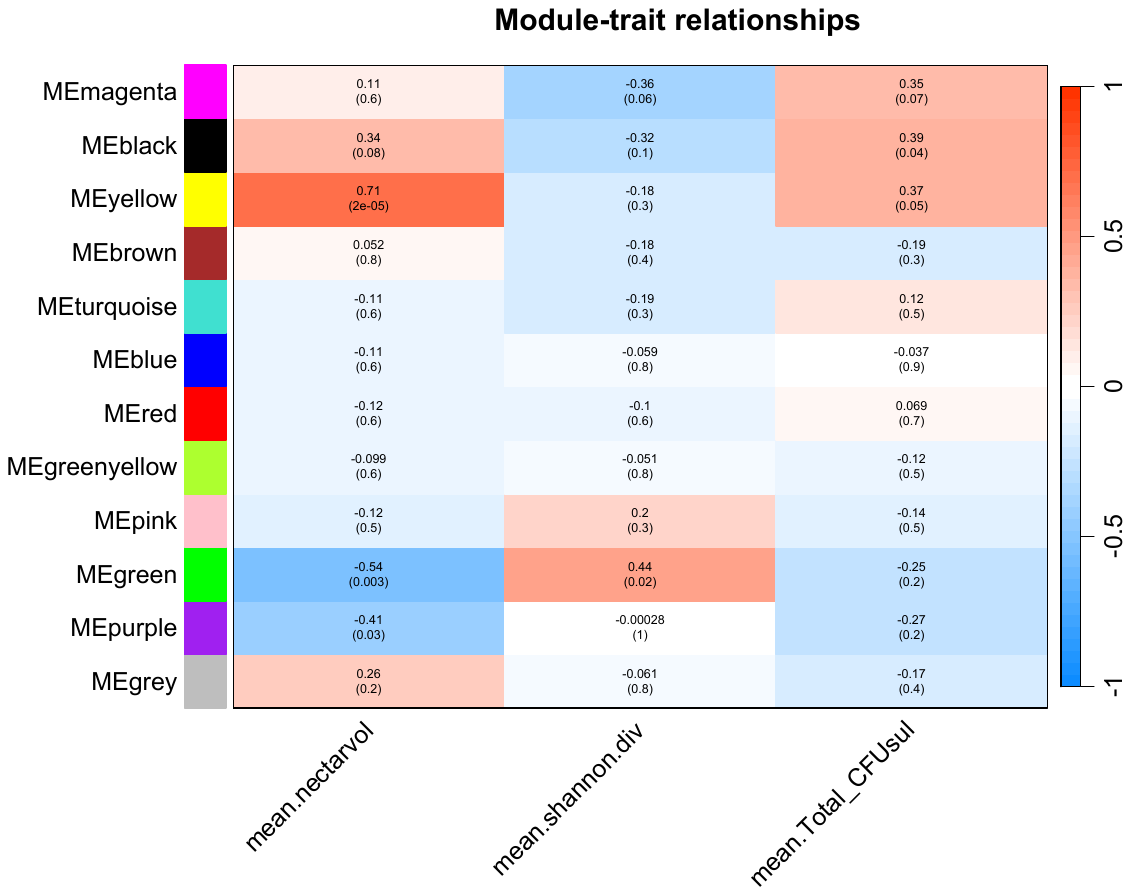
